# Deposition of Polycyclic Aromatic Hydrocarbons (PAHs) into Northern Ontario Lake Sediments

**DOI:** 10.1101/786913

**Authors:** Graham A. Colby

## Abstract

Polycyclic aromatic hydrocarbons (PAHs) are priority pollutants that are produced through incomplete combustion of modern biomass and fossil fuels. In aquatic systems PAHs are absorbed by suspended matter and ultimately deposited into sediments. PAH fluxes to sediments have been declining in North American since the mid 1960s. Improving technology and regulations were expected to contribute to declining PAH concentrations; however, in some urban sediment there are recent increases in deposition. Trends in concentrations of pyrogenic PAHs and perylene were determined in the sediment of two lakes, in central Ontario. Intact piston cores that preserve the depositional history were collected from each site, sliced into 1 cm intervals and analyzed using gas-chromatography/ mass-spectrometry. Pyrogenic PAH trends at each site displayed unique characteristics suggesting differing extents of influence from various atmospheric sources. The upper core profile (above 8.5 cm) in the more remote site had decreasing PAH concentrations consistent with observations from Siskitwit Lake. The more urban site (above 3.5 cm) had increasing PAH concentrations suggesting modern anthropogenic activities have a larger influence in this region. Perylene fluxes at both sites do not correlate with the observed PAH fluxes, increasing in concentration with depth, thus indicating separate sources for this PAH, likely diagenesis within the sediments. Both sites had PAH concentrations exceeding the interim sediment quality guidelines in the uppermost sediment deposits. This study provides insights into the differential atmospheric deposition in Ontario and may aid in establishing strategies for reducing or mitigating the production of PAHs.

## 1.0 Introduction

Polycyclic aromatic hydrocarbons (PAHs) are a complex group of organic chemicals with a fused ring structure of at least two benzene rings in linear, angular or cluster arrangements. PAHs are released into the environment as products of incomplete combustion from the burning of fossil fuels (eg., power generation and transportation) and other anthropogenic processes such as manufacturing (eg., aluminum and coke production). PAHs are persistent organic pollutants (POPs) of increasing concern because of their ubiquity in ambient air and carcinogenic properties (Nielsen et al., 1996; Bofetta et al., 1997). These environmental pollutants accounted for an estimated 5 million tons of toxicants released in the US in 2000 (Yan et al., 2003). Epidemiologic data indicates a relationship between occupational and environmental exposure to PAHs and increased risk of cancer (Bofetta et al., 1997). The environmental abundance of PAHs can present a threat to both human and environmental health (Bofetta et al., 1997; Yan et al., 2003). Several PAHs, notably benzo[a]pyrene, benz[a]anthracene, chrysene, benzo[b]fluoranthene, benzo[j]fluoranthene, benzo[k]fluoranthene, and ideno[1,2,3-cd]pyrene are linked to health concerns because of their strong carcinogenic properties (Ravindra et al., 2008a). Industrialized and urban centers frequently have more degraded air and sediment quality associated with increased energy generation and vehicle traffic (Mastral and Calen, 2000; Van Metre et al., 2000). The majority of the PAHs produced are deposited within a kilometer of the source; however, a fraction can be transported thousands of kilometers before deposition (Lima et al., 2005). Due to the widespread sources and environmental persistence, PAHs disperse globally through atmospheric transport (Bjorseth et al., 1979; Wilcke, 2007; Lee and Vu, 2011). This study aims to understand the historical fluxes of PAH deposition in central Ontario. Given that Canada’s energy consumption is projected to increase 34% by 2025 (Hofman and Li, 2009), with the leading component being oil, it is more pertinent than ever to understand the environmental impact of our energy requirements. As demonstrated in this study atmospheric deposition of PAHs has the potential deposit contaminant in excess of sediment quality guidelines defined by the Canadian Council of Minister of the Environment. The results presented may have an integral role in: allocating responsibility for the contaminant concentrations, and establishing mitigation and remediation policies.

### 1.1 Priority PAHs

The United States Environmental Protection Agency (EPA) has defined 16 PAHs as priority pollutants (Yan at al., 2003). Although PAHs often occur in complex mixtures of differing environmental toxicity these 16 compounds were selected as priority pollutants because of i) toxicity ii) extent of available information, iii) the likelihood of human exposure and iv) environmental occurrences (ATSDR, 1995). Each of these 16 PAH compounds have unique chemical and toxicological properties that effect their environmental availability and abundance (Ravindra et al., 2008a).

### 1.2 Sources and Formation

PAHs found in the environment form by one of three possible processes: pyrogenically, petrogenically, or diagenetically. The most prominent and ubiquitous source in the environment is pyrogenic PAHs synthesized from: high temperature, short duration, incomplete combustion of organic matter (Meyer and Ishiwatari, 1993).

During pyrogenisis organic compounds are partially cracked to lower molecular weight radicals and unstable compounds (Douben, 2003). This occurs when reaction temperatures exceed 500°C, providing sufficient energy to break carbon-hydrogen and carbon-carbon bonds, forming free radicals (Ravindra et al., 2008a). These fragments, mainly highly reactive free radicals, lead to more stable aromatic formations through pyrosynthesis reactions (Ravindra et al., 2008a; Mastral and Callen, 2000). High molecular weight PAHs (∼500-1000 amu) function as precursors to soot formation (Lima et al., 2005; Ritcher and Howard, 2000). PAHs and soot carbon molecules form strong interactions from formation through to deposition in the aquatic environment, significantly affecting PAH partitioning and availability (Douben, 2003).

Petrogenic PAHs are a natural component of unburned petroleum and can be released directly to the environment through human or natural processes (eg. crude oil spills). Petrogenic PAHs occur naturally through digenetic processes over geological time scales, resulting in the formation of petroleum products and fossil fuels (Boehm et al. 2001). Petrogenic PAHs form at relatively low temperatures (150°C) over long time periods and are highly alkylated, reflecting the ancient plant parent material (Douben, 2003). The alkylated PAH content is a characteristic feature used in the identification of petrogenic PAH sources (Wang et al., 1999).

The third potential method of production is biotic diagenetic reactions. Diagenetic PAHs refer to those formed from biogenic precursors such as plant terpenes leading to the formation of certain precursor molecules that can yield perylene, phenanthrene and chrysene derivatives (LaFlamme and Hites, 1979; Hites et al., 1980; Douben, 2003). The specific reaction mechanism remains unknown, however it is thought to involve an anaerobic process (Venkatesan, 1988). Increasing concentrations of these compounds with increasing depth of the sediment core is often interpreted as in-situ diagenesis (Hites et al., 1980; Wakeham et al., 1980; Slater et al., 2013). Although diagenetic PAHs appear as background levels in recent sediments (deposited over the last 100 years) they dominate the profile in older sediments prior to industrial activity (Gschwend et al. 1983).

#### 1.2.1 PAH formation from combustion reactions

Type of fuel, amount of oxygen, and temperature are factors that affect the production and environmental fate of combustion-derived PAHs. The variations in these three factors make it difficult to predict the relative proportions of pyrogenic PAHs.

The type of fuel burned influences the growth mechanism of PAHs and the amount released during combustion. Experiments have indicated that increasing the amount of aromatic content in fuel elevates the PAH emissions (Ritcher and Howard, 2000; Lima et. al, 2005). This trend is linear for many PAHs (Lima et al., 2005); however, assortments of PAHs are emitted regardless of their presence in the original fuel. The relative PAH proportions in emissions do not necessarily coincide the original fuel, as PAHs originally present are broken down and reformed.

Formation of pyrogenic PAHs requires an oxygen-depleted environment. Increasing the amount of excess oxygen leads to a more efficient combustion process to the point of complete combustion. Studies have shown that increasing the amount of airflow reduces PAH concentrations and soot formation (Lima et al., 2005). The amount of available oxygen for combustion can be described in terms of the air relative to fuel ratio (A/F). High A/F supplies larger quantities of oxygen and more efficient combustion can occur resulting in lower PAH emissions.

Temperature has also been correlated with the molecular distribution of PAHs. Low combustion temperatures generate enriched alkyl-substituted PAHs, in contrast higher combustion temperatures that produce more parent-like PAH compounds (LaFlamme and Hites, 1978).

#### 1.2.2 Sources

As there are no identifiable petrogenic sources in the systems studied, pyrogenic PAHs produced through combustion and perylene produced diagenetically are the subject of this study. Anthropogenic emissions from vehicles, heating, power plants, industrial processes and open burning are considered the principal PAH sources (Baek et. al, 1991; Van Metre et al., 2000). Pyrogenic PAHs enter the environment through three major emission sources: domestic, mobile, and industrial.

Domestic emissions are predominantly associated with the burning of coal, oil, garbage, gas and other organic matter for heating and cooking purposes (Lee and Vu, 2011). The low-temperature combustion of these fuels has the potential to result in higher emissions of PAHs than high-temperature industrial combustion sources (Ravindra et al., 2008a). Domestic sources are consequently important contributors to the total emissions of PAHs in the environment. Research has indicated a strong correlation between urbanization and population density with the overall abundance of PAHs (Van Metre et al., 2000; Hafner et al., 2005).

Mobile emissions from vehicles such as aircraft, shipping, railways and automobiles are a major source PAHs found in urban and remote areas. Studies have shown that emissions from vehicle exhaust are the largest contributors of PAHs in urban areas (Shen et al., 2011). As mentioned previously the production of PAHs from gasoline in automobiles depends on the air/fuel ratio. Along with improving engine air/fuel ratio, catalytic converters on automobiles have significantly reduced the PAH exhaust emissions (Ravindra et al., 2008a). As a result of improving technologies and robust regulations the relative contribution of motor vehicles to total PAH emissions is expected to decrease, reducing human exposure and potentially accounting for decreasing PAH fluxes observed in the upper sediment layers (Hites et al., 1977; VanMetre and Mahler, 2005; Shen et al., 2011).

Industrial sources include: primary aluminum production, coke production, creostote and wood preservation, waste incineration, cement manufacturing, petrochemical industries and related commercial heat/power production (Government of Canada, 1994). Although these emissions are released from stationary sources they enter the atmosphere remaining airborne for days and impacting regions outside of their immediate vicinity, thousands of kilometers away (Ravindra et al., 2008b). Air pollution control devices on industrial stacks minimize the harmful release of PAHs having shown to remove high molecular weight PAHs with four or more rings with 78% efficiency and low molecular weight PAHs with two or three rings with 5% efficiency (Lee and Vu, 2011).

Comparatively natural sources have a smaller overall contribution to environmental PAH content and are considered negligible. Natural sources of PAH include those of terrestrial origin (eg. volcanoes and non-anthropogenic burning of forest) and cosmic origin (eg. chondrites). Natural PAHs are produced through: high temperature synthesis, low temperature diagenesis of organics to fossil fuels, and direct biosynthesis by microbes and plants (Ravindra et al., 2008a).

### 1.3 PAH Properties

The transport, deposition and chemical transformation of PAHs is strongly dependent on their respective gas/particulate phase partitioning, vapor pressure, molecular weight and environmental conditions including temperature, humidity, precipitation and type of particles in the atmosphere (Harner and Bidleman, 1988; Baek et al., 1991; Lima et al., 2005). Low molecular weight (LMW) PAHs containing 2-3 aromatic rings tend to be emitted in the gaseous phase, while high molecular weight (HMW) PAHs of 4 or more aromatic rings tend to be emitted in the particulate phase (Lee & Vu, 2011). With the exception of a few lighter compounds (naphthalene, acenaphthene, acenaphthylene, and fluorene) most PAHs have low vapour pressures, are relatively non-volatile and have a low solubility in water. The solubility and vapor pressure of PAHs generally decrease with increasing molecular weight, thus smaller PAHs are generally more volatile than larger PAHs.

The structure and physical properties control PAH volatility, solubility, sorption, and decomposition behaviours (Lima et al., 2005). Water solubility values range from highly insoluble (benzo[ghi]perylene; 0.003 mg/L at 25 ° C) to slightly soluble (naphthalene; 31 mg/L at 25 ° C) and vapor pressures range from highly volatile (naphthalene; 10.4 Pa at 25°C) to relatively non-volatile (dibenzo[a,h]anthracene; 3.70^−10^ Pa at 25°C) (Appendix A Table A1; Douben, 2003).

### 1.4 PAH Distribution

PAHs are stable and persistent hydrocarbons, leading to their wide distribution in sediments throughout the world, with their abundance increasing with proximity to urban centers (LaFlamme and Hites, 1978; Hafner et al., 2005). Hafner et al. (2005) observed that sites within 25 km of coastal regions have lower than expected atmospheric PAH concentration based on population size due to dilution with cleaner air, while sites near industrial outputs or point sources have higher than expected concentrations. Additionally Hafner et al. (2005) noted significantly higher atmospheric PAH concentrations in developing countries compared to the rest of the world, indicating that regulations and technological innovations, particularly the substitution of coal with cleaner burning fuels successfully reduce PAH pollution.

PAHs found in the gas phase have relatively short atmospheric lifetimes, on the order of hours to days (Lima et al., 2005). Larger 5-6 ring PAHs deposit slowly depending on atmospheric conditions, may be airborne for days, and transported over longer distances. A study conducted at various sites in Belgium found vapour phase PAHs shared more source-specific characteristics than aerosol phase PAHs (Ravindra et al., 2008b). Most combustion-derived PAHs are associated with conglomerate particles such as Black Carbon and soot. The residence times for particulate PAHs are comparatively longer spending four or more days in atmospheric suspension (Ravindra et al., 2008b). Particulate deposition, just like individual PAH deposition, varies with size. Windsor and Hites (1979) determined small particles (1 μm in diameter) are transported for about 1300 km before deposition, while large particles (10 μm in diameter) settle much closer to the source at about 13 km (particle density = 2g/m, height = 20 m and wind velocity = 4m/s).

Atmospheric PAHs are found mostly in particulate form. Once in the atmosphere PAHs can be: i) removed by wet or dry deposition, ii) transported through shifts in air masses, turbulence and convection, iii) degraded (or transformed) by chemical and photochemical mechanisms, and iv) exchanged between gas and particle phases based on equilibrium shifts (Baek et. al, 1991). Dry and wet deposition from the atmosphere depends on the compound’s physiochemical properties (solubility and vapor pressure) and meteorological parameters (rain volume, intensity and temperature) (Ravindra et al., 2008a). In general PAHs in the gas phase dissolve in raindrops and are subject to wet deposition (Baek et. al, 1991; Ravindra et al., 2008a). PAHs bound to particles are subject to both wet and dry deposition. The rain can directly wash them out; however, larger particles also experience direct fallout dependent on their size (Windsor and Hites, 1979; Lima et al., 2005). PAH deposition onto land will slowly be degraded by microbial activity, while deposition onto aquatic systems will cause the hydrophobic PAHs to aggregate into conglomerate particles. These larger conglomerates are forced out of the aqueous phase to reduce the entropic cost associated with hydrating each particle, finally settling to the sediment floor. Sediments are the main environmental sink for PAHs, as PAHs within sediments experience minimal photochemical or biological degradation and are well preserved (Douben, 2003).

### 1.5 Source Identification

PAHs are emitted to the environment from a variety of sources, causing uncertainty when delineating the specific influential production source(s). Efforts to determine the specific origin is challenging because atmospheric deposition of PAHs is likely not the result of one industry or activity. However, source apportionment is a crucial exercise to determine control strategies for environmental pollutants. Several propitious methods have been used to apportion PAH sources including: source markers (profile based), and diagnostic ratios.

Source (or profile) markers rely on the concept that specific PAHs indicate certain processes. The concentration profile can be used to determine the contribution of different sources to the concentration in the air (Khalili et al., 1995; Simcik et al., 1999). Khalili et al. (1995) determined the chemical composition of PAHs from airborne sources and found that the abundance of 2-3 ring PAHs differed significantly depending on the source (Ravindra et al., 2008a). Additionally, 3 ring PAHs chrysene and benzo[b,k]fluoranthene production have been associated with coal combustion (Khalili et al., 1995; Ravindra et al., 2008a). These source markers are often preserved in sediment profiles.

Diagnostic ratios compare ratios of frequently identified PAHs. The benz[a]anthracene/(benz[a]anthracene + chrysene/triphenylene) [BaA/ Σ 228]; fluoranthene/(fluoranthene + pyrene) [Fl/(Fl+Py)]; and indeno(1,2,3-cd)pyrene/(indeno(1,2,3-cd)pyrene + benzo(g,h,i)perylene) [IP/(IP+Bghi)] ratios are applied extensively to differentiate between pyrogenic and petrogenic PAH sources (Yunker et al., 2002; Lima et al., 2005). The distribution of alkylated versus parent PAHs in sedimentary environments is also used to distinguish between high and low temperature sources (Youngblood and Blumer, 1975).

The validity of these approaches has been questioned as they rely on the assumptions that: i) each suspected source or source type has relative proportions of PAH species that are unique, and ii) that the relative proportions of the species remain conserved between emission source and point of measurement (Galarneau, 2008). Source markers and ratio based source apportionment methods compare the source and sink data without accounting for alterations that may occur in the atmosphere. One issue that potentially arises in these assumptions has been previously mentioned: fuel type, and air/fuel affect the PAHs produced. PAH source signatures are not entirely unique to each source type. With regards to the second assumption, due to volatility, reactivity and partitioning, the relative PAH concentration is not conserved in the atmosphere (Galarneau, 2008). These assumptions hold true only under limited conditions and utilization of the other apportionment methods are necessary to more accurately identify the potential sources and correct for errors resulting from atmospheric processes.

### 1.6 Sediment Profiles

The use of lake sediments to identify the onset and variation in current and past environmental contamination is well established. Numerous studies have analyzed the PAH flux in lake sediments (Hites et al., 1977; LaFlemme and Hites, 1978; Hites et al., 1980; Fernadez et al., 2000; Kannan et al., 2005). Present day effects of airborne contamination can be placed in a historical context using sediment profiles. In the absence of long-term monitoring, lake sediments serve as a reliable means to analyze past contamination. PAHs are hydrophobic and hence have a higher tendency to associate with particles than to dissolve in water (represented by the octanol-water partition coefficients; Appendix A Table A1). In aquatic systems PAHs have an affinity for settling particles (Figure 1.6). The strong absorption of PAHs onto particles (particularly Black Carbon and soot) can reduce bioavailability slowing degradation rates (Douben, 2003; Lima et al., 2005). With the use of dating methodologies an intact core can accurately preserve a historical record of PAH concentration.

The PAHs found in sediments often differ in content from those found in petroleum, supporting the conclusion that most PAHs in aquatic environments derive from pyrogenic sources (Hites et al., 1977; Douben, 2003). The dominance of parental PAHs over alkyl-substituted homologs in sediment profiles and lack of a distinctive unresolved complex mixtures precludes petroleum sources as contributors to sedimentary PAH profiles (Youngblood and Blumer, 1975; Wang et al. 1999). Petrogenic PAHs are generally associated with local or point sources; in contrast pyrogenic PAHs are distributed on a global scale (Hites et al., 1977). Pyrogenic PAHs are more widely distributed, more persistent and largely protected from photochemical and microbial degradation because of their adherence to soot carbon (Douben, 2003). When petrogenic PAHs (crude oil or petroleum products) are accidentally released to the environment they are subject to a variety of weather processes including: evaporation, dissolution, microbial degradation, and other dispersive processes such as photoxaidation and adsorption to suspended material (Wang and Fingas, 2003).

Trends in PAH deposition over time can be deduced through analysis of PAH concentrations in lake sediments. Atmospheric deposition is the prevailing source of PAHs to lakes, so trends in the core indicate the long-term record changes in the atmospheric loading of these pollutants. Lead-210 dating can be used to determine the rate of sedimentation, providing a timeframe for deposition and the ability to calculate fluxes in PAH deposition (Robbins and Edgington, 1975; Evans & Rigler, 1982). To accurately identify historical pyrogenic combustion suitable lakes should: i) be relatively removed from point sources, ii) receive PAHs mostly via atmospheric deposition, iii) have small catchment basins to minimize PAH residence time, iv) have rapid sediment accumulation and v) have minimal postdepositional mixing (ie. lack of bioturbation) (Lima et al., 2003). Ultimately this allows for the preservation of vertically expanded, highly accurate historical contamination records.

Sedimentary records show a strong correlation between PAH concentration profiles and energy consumption associated with industrialization. A typical PAH profile reveals a gradual increase in total PAH concentrations beginning around 1880, with the onset of the Industrial Revolution, to a maximum in the 1950s (Lima et al., 2003). The substitution of coal with cleaner-burning fuels, stricter emission policies, and the use of catalytic converters explains the decrease in PAH concentrations from the 1960s onward (Lima et al., 2005). Despite this declining trend noted in many studies (McVeety and Hites, 1988; Slater et al., 2013), other recent studies have indicated the total PAH concentrations are increasing in certain areas of the United States (Van Metre and Mahler, 2005). This increase in PAHs parallels increased use in automobile usages implying a strong correlation between PAH deposition and urban sprawl.

## 2.0 Research Scope

The primary objective of this study is to build upon previous literature and to assess if atmospheric deposition of pyrogenic PAHs is occurring in lakes in central Ontario. Previous findings suggest that PAH concentrations decrease in the upper sediment level compared to the high concentrations reflective of early industrial activity at the bottom of the profile (Hites et al., 1977; Wakeham and Schaffner, 1979; Van Metre et al., 2000). This study aims to verify these findings in lakes from central Ontario and to compare the observations to previously studied sediments from Siskiwit Lake to determine whether there are regional or local controls. The proximate research question will elucidate i) which PAHs are present in the lake sediments, ii) what are their relative proportions and iii) how the deposition at these sites compare with other findings in regional lakes. The expanded research question will assess the contribution PAHs from regional and local sources. Through a comparison of the complex PAH profiles of these sites with existing literature we will investigate whether these contaminants were deposited atmospherically. These findings have environmental implications that will contribute to potential remediation and mitigation efforts.

We hypothesize that the PAHs implicated in this study will be the result of pyrogenic atmospheric deposition, and will therefore consist of a larger proportion of high molecular weight PAHs. In comparison to other sites the studied lakes may share a general overlaps based on shared regional sources, or similar contributing sources; however, if local sources that are specific to each lake contribute to the contaminant profiles, their impact will likely exceed the regional sources due to proximity between the source and the sink.

## 3.0 Materials and Methods

### 3.1 Study Area and Sediment Sampling

Solitaire Lake is located in central Ontario just outside of Huntsville, while Fairbank Lake is part of the Greater Sudbury Region situated approximately 221 km North West of Solitaire Lake (Figure 3.1). Both of these lakes are partially surrounded by Provincial Parks – J. Albert Bauer and Fairbank Provincial Park, respectively, and the other adjacent properties are private cottages. With the exception of limited pleasure craft operation the study areas lack significant anthropogenic point sources. Consistent with previous studies it is assumed that all pyrogenic PAH sediment loadings are derived from atmospheric deposition. Additionally both of these sites have small catchment areas, which minimizes the residence time of the PAHs and reduces postdepositional mixing, allowing for the preservation of vertical historical records.

**Figure 3.1:**
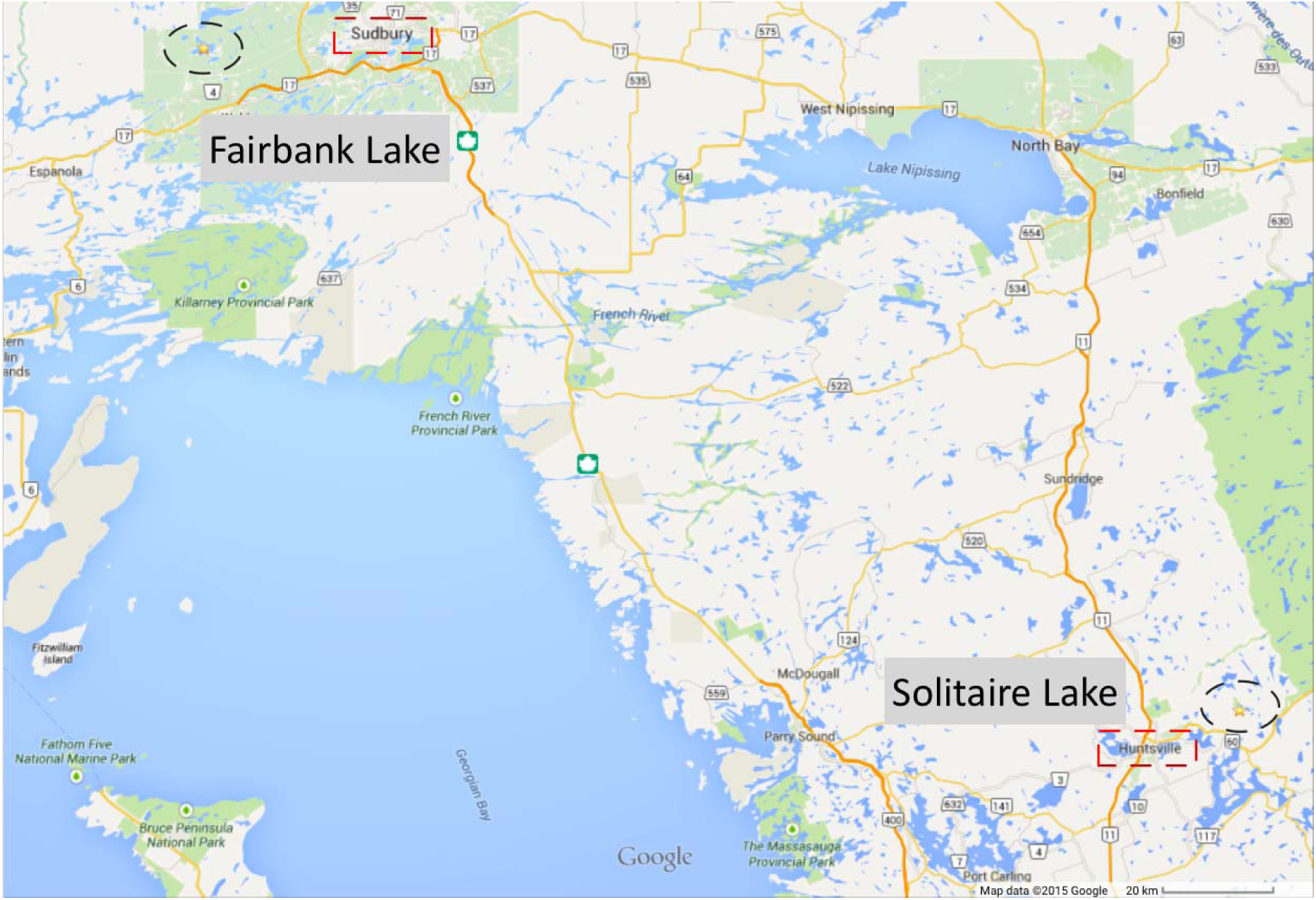
Sample locations (GoogleMaps, 2015). Fairbank Lake, is located west of Sudbury indicated by the star. Solitaire Lake, is located east of Huntsville and indicated by the star.

Solitaire L. was sampled at 45°23’22.7 N, 79°00’39.3 W (DMS) at a depth of 98.2 ft. Fairbank L. was sampled at 46°27’33.5 N, 81°25’53.3 W (DMS) at a depth of 149 ft. Bathymetry data was accessible for Fairbank L. (Appendix B Figure 1), while we relied on local knowledge to determine the maximum depth of the basin in Solitaire L. The expected depth measurements were confirmed with a Hummingbird© fish finder and the GPS coordinates were captured using a Garmin© eTrex20. Five-piston cores 6.5 cm in diameter were collected from the deepest region of each lake using a gravity corer, with two additional 5 kg weights attached. It is assumed the coring device collected vertically stratified sediments, with minimal compaction and mixing upon impact and removal. The cores ranged between 30-60cm in length, and multiple cores were collected from approximately the same location at each lake. The cores were sliced into 1 cm increments on shore after removal, placed in Whirl-Pak© bags, stored in the dark, and kept cool (with ice and dry ice) until they could be returned to the lab. Once at the lab the samples were frozen at −20°C until further processing.

### 3.2 Standard Preparation and Quality Control

Before sample extraction and clean up, standards of known PAHs were prepared for compound identification, quantification and quality control. A working standard of SV Calibration Mix #5 (Restek; Cat: 31011; Lot: A064861) PAH reference standard (SV Calibration Mix #5) containing the 16 priority PAHs and perylene (Sigma; Cat: P11204-1G; Lot: 04215DY) were prepared to stock concentrations of 1, 3, 5, 10, 15, 25, 35, 45, 80 and 100 ppm. The standard curves had a linear correlation for all PAHs between 3-80 ppm. o-terphenyl (Sigma; Cat: T2800-25G; Lot: 14530TA) was used as an internal standard. 9,10-dihydrophenanthrene (Sigma; Cat: D106003-5G; Lot 12812JC) and m-terphenyl (Sigma; Cat: T3009-5G; Lot: 12514CA) were used as recovery standards. The standards were selected because they are not found in natural systems, and have retention times that are distributed between the retention times of the 16 priority PAHs. The internal standard was prepared to 121.77 μg/mL adding 10 μL to all samples. The recovery standards were prepared to 128.7 μg/ml (for m-terphenyl) and 119.38 μg/ml (for 9,10-dihydrophenanthrene) stock solutions injecting 15 μL into each sample immediately prior to microwave extraction.

Quality control was maintained extracting 1 processed blank per core. The accuracy of the method was determined using 2 SV Calibration Mix #5 standards and 2 EC-1 (Environment Canada, reference material) per whole lake core. The processed blanks had no detectable PAHs and were free of peaks above the column background. The average recovery of each PAH in the solvent spiked with SV Calibration Mix #5 was greater than 75%±10, with an 80% recovery for naphthalene. The average recoveries for compounds identified in the EC-1 reference material were 108%±20. External standards (9,10-dihydrophenanthrene and m-terphenyl) recoveries varied between Solitaire L. and Fairbank L. GC/MS runs. In the Solitaire L. analysis the 9,10-dihydrophenanthrene recoveries were on average 80%+30 and the m-terphenyl recoveries were on average 90%±30. In the Fairbank L. analysis the 9,10-dihydrophenanthrene recoveries were on average 70%±10 and the m-terphenyl recoveries were on average 80%±10. The relative standard deviation (RSD), used in the result plots for each PAH in the sediment core, was measured by triplicate o-terphenyl standard injections that were applied to each section within that core (± 20% in Solitaire L. and ± 10% in Fairbank L.). The PAH concentrations in each sample were corrected for using the 9,10-dihydrophenanthrene recovery to correct PAHs with molecular ion masses between 128-178 m/z, and m-terphenyl to correct PAHs with molecular ion masses of 202-278 m/z. The limit of quantification for parent PAH compounds was 12.5 ng/g dw in Solitaire L. samples and 8.5 ng/g dw in Fairbank L. samples. The difference in the limit of quantification results from different average dry weights between the two sites.

### 3.3 Sample Preparation

Each sample was freeze-dried recording wet mass and dry mass, from which wet and dry bulk density was calculated (Appendix B Table 1A-C). The samples were then stored in the dark until extractions could be completed. PAH analysis was performed using modified EPA methods 8100, 3051A, and 3630C (Slater et al., 2013). The Fairbank L. core had high dry masses in each core section (>4 g) and was sufficient to analyze on its own, while the Solitaire L. core sections had low dry masses in each core section (<3 g) and the two most similar cores in depth and bulk density were combined to gain better detection. The sufficient mass for analysis was about 2 g dw or more from each section (1 cm interval) of the core. Approximately 3-8 g of sediment were extracted per core section. All samples and standards were spiked with 15 μg/mL of external recovery standards (9,10-dihydrophenanthrene at 1410 ug/ml and m-terphenyl at 1267 ug/ml), then microwave extracted using a MarsX© in 40-45 mL of 1:1 hexane/acetone at 115°C for 10 minutes. Extracts were allowed to cool to approximately 30°C and then filtered with approximately 80 mL of hexane through Whatman© glass fiber filter paper. Filtered extracts were reduced in volume to approximately 1 mL, before a copper cleanup was completed to remove sulfur compounds. Copper pellets were acid rinsed with 30% nitric acid before rinsing 3 times each with methanonl (MeOH), dichloromethane (DCM), and hexane. The copper pellets were dried under nitrogen, before being added to each sample. The samples settled for at least 4 hours before the hexane fraction was removed and the vial was rinsed 3 times with approximately 0.5 mL of hexane.

The samples were then blown down to 1 mL before loading onto the activated silica gel chromatography column. The samples were loaded onto 7 g of activated silica gel in hexane. Four fractions were collected: F1 hexane (aliphatic hydrocarbons); F2 2:1 hexane/DCM (aromatic hydrocarbons); F3 DCM; F4 methanol. F3, and F4 were archived while the F1 and F2 fraction were reduced to 100 μL volume under nitrogen, solvent exchanged into hexane (for the case of F1), and spiked with 5 μg/mL internal standard (o-terphenyl). Napthalene was present in both the F1 and F2 fractions while all other identified PAHs were only present in the F2 fraction. The F1 and F2 fractions were processed, and concentrated to a total volume of 100 μL, injecting 2 μL onto the GC/MS column.

### 3.4 Gas Chromatography-Mass Spectrometry (GC/MS)

Using micro-inserts, the samples were placed in 1 mL amber GC vials for sample analysis by GC/MS. PAH quantification was performed on GC/MS (Agilent 6890 GC, Agilent 5973n MS) equipped with a split/splitless injector kept in splitless mode and a DB5-MS capillary GC column (with a 30 m length x 0.32 diameter and 0.25 *μ*m film thickness). The oven was programmed to 50°C (5 minute hold) and ramped at 4°C/min to 320°C (held for 12 min). The flow of helium gas was 1.2 mL/min. A total scan of ions 50-450 MW was completed. The following 16 EPA priority PAHs were quantified in addition to perylene: naphthalene, acenapthylene, acenaphthene, fluorene, phenanthrene, anthracene, fluoranthene, pyrene, benz[a]anthracene, chrysene, benzo[b]fluoranthene, benzo[k]fluoranthene, benzo[a]pyrene, benzo[g,h,i]perylene, dibenz[a,h]anthracene, and ideno[1,2,3-cd]pyrene. All of the PAHs were quantified using single ion extractions (Appendix B Figure 2A-B, Appendix B Table B2) with at least 7-point calibration curves, prepared relative to the internal standard.

## 4.0 Results

### 4.1 Total Pyrogenic PAH Concentrations

The total pyrogenic PAH concentrations (exclusive of perylene and unconfirmed benzo[e]pyrene) in ng/g dw are plotted versus depth in cm for Solitaire L. (Figure 4.1.1) and Fairbanks L. (Figure 4.1.2). PAH concentrations were corrected for using the recovery standards, and will be the basis of further discussion throughout. Notably the correction for the recoveries does not change the overall trends in the data for either site.

**Figure 4.1.1:**
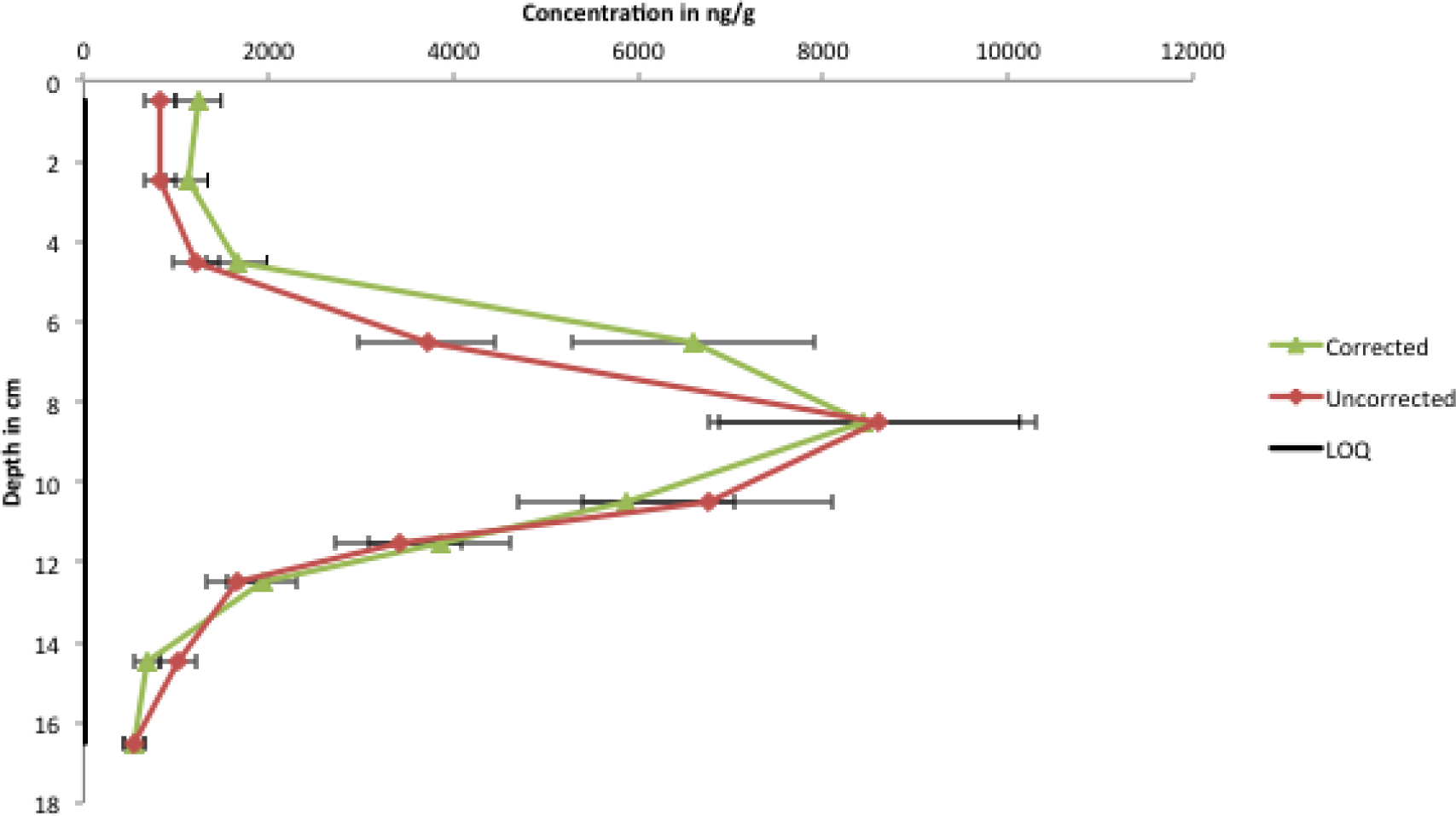
Total PAH concentration for Solitaire L. The figure indicates: the concentrations corrected for the losses in method using the recovery standards and the uncorrected concentrations. The horizontal error is a 20% relative standard deviation based on o-terphenyl standard replicates. The vertical black line indicates the limit of quantification (LOQ).

**Figure 4.1.2:**
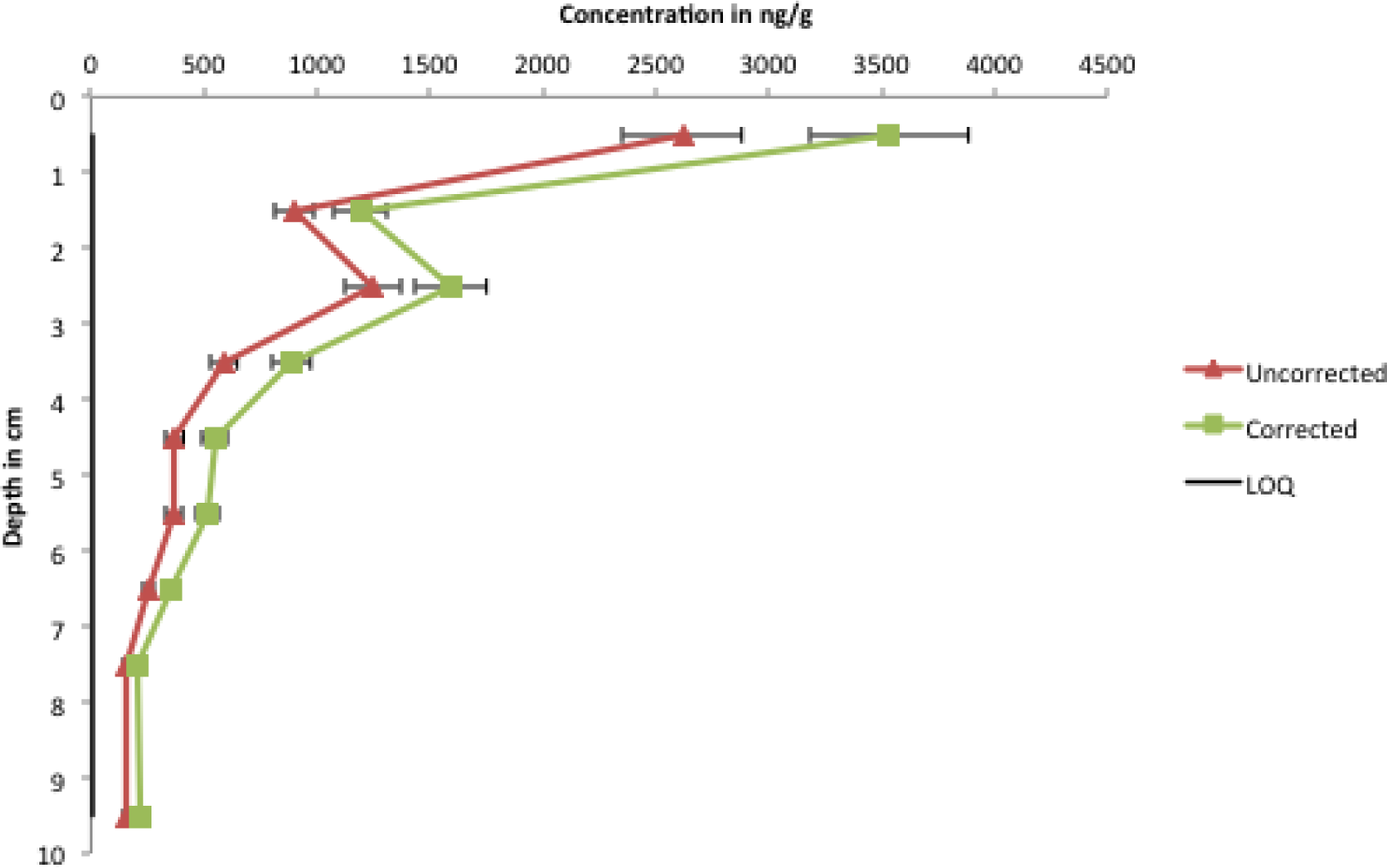
Total PAH concentration for Fairbank L. The figure indicates: the concentrations corrected for the losses in method using the recovery standards and the uncorrected concentrations. The horizontal error is a 10% relative standard deviation based on o-terphenyl standard replicates. The vertical black line indicates the limit of quantification (LOQ).

The total PAH concentrations each have distinguishing features. In Solitaire L. the pattern of PAH fluxes followed those observed in earlier studies (McVeety and Hites, 1988; Slater et al., 2013). The profile exhibits lower concentrations of at the top and bottom of the sediment profile, with a maximum peak of approximately 8500 ng/g dw occurring at approximately 8.5 cm. The total PAH concentration was approximately 600 ng/g dw or less at the lower depths (below 14 cm), and approximately 1500 ng/g dw or less at the upper depths (above 5 cm).

In Fairbank L., the pattern of PAH fluxes followed trends observed in recent studies (Lima et al., 2003; VanMetre and Mahler, 2005). The profile exhibits lower concentrations at the bottom of the sediment profile, with a relative maximum peak of approximately 1500 ng/g dw occurring at approximately 2.5 cm, and then increasing to approximately 3500 ng/g at the uppermost portion of the profile. The total PAH concentration was approximately 300 ng/g dw or less at the lower depths (below 4 cm), and approximately 1000 ng/g dw or more at the upper depths (above 2 cm).

Assuming the middle peaks in each profile (8.5 cm in Solitaire L. and 2.5 cm in Fairbank L.) are consistent with the mid to late 1950s peak observed by McVeety and Hites (1988), it can be noted that Fairbank L. has a higher sedimentation rate leading to a more compact profile. This is also noticeable in the dry mass per cm^3^ and dry bulk density, which were significantly larger in Fairbank L., justifying the necessity of combine two Solitaire L. cores (Appendix B Table B1A-C).

### 4.2 Relative Abundance

Thirteen of the seventeen PAHs identified (excluding naphthalene, fluorene and perylene) exhibit a similar trend within the sites (Figure 4.2.1 and 4.2.2). In both locations naphthalene, fluorene and perylene were inconsistent with the overall trend in total PAH concentration. The variation in both naphthalene and fluorene at both sites can likely be attributed to high volatility of these two low molecular weight compounds as well as the relatively low concentrations at which they occur within the sediment, resulting in poor detection. The naphthalene profiles are aberrant with a distinct maximum peak 2 cm above in Solitaire L. and 2 cm below in Fairbank L. in comparison to other pyrogenic PAH peaks, suggesting a different source for this compound in the profile. In Solitaire L., the perylene profile peaks at 6.5 cm, 2 cm above the total PAH profile. Intriguingly, the perylene profile indicates increasing concentrations in both the upper and lower sections of the core, where all other quantified PAHs are decreasing. In Fairbank L., the top segment of the profile (above 3 cm) corresponds with the observed total PAH profile; however, below 3 cm the perylene profile deviates indicating an increasing concentration with depth.

**Figure 4.2.1:**
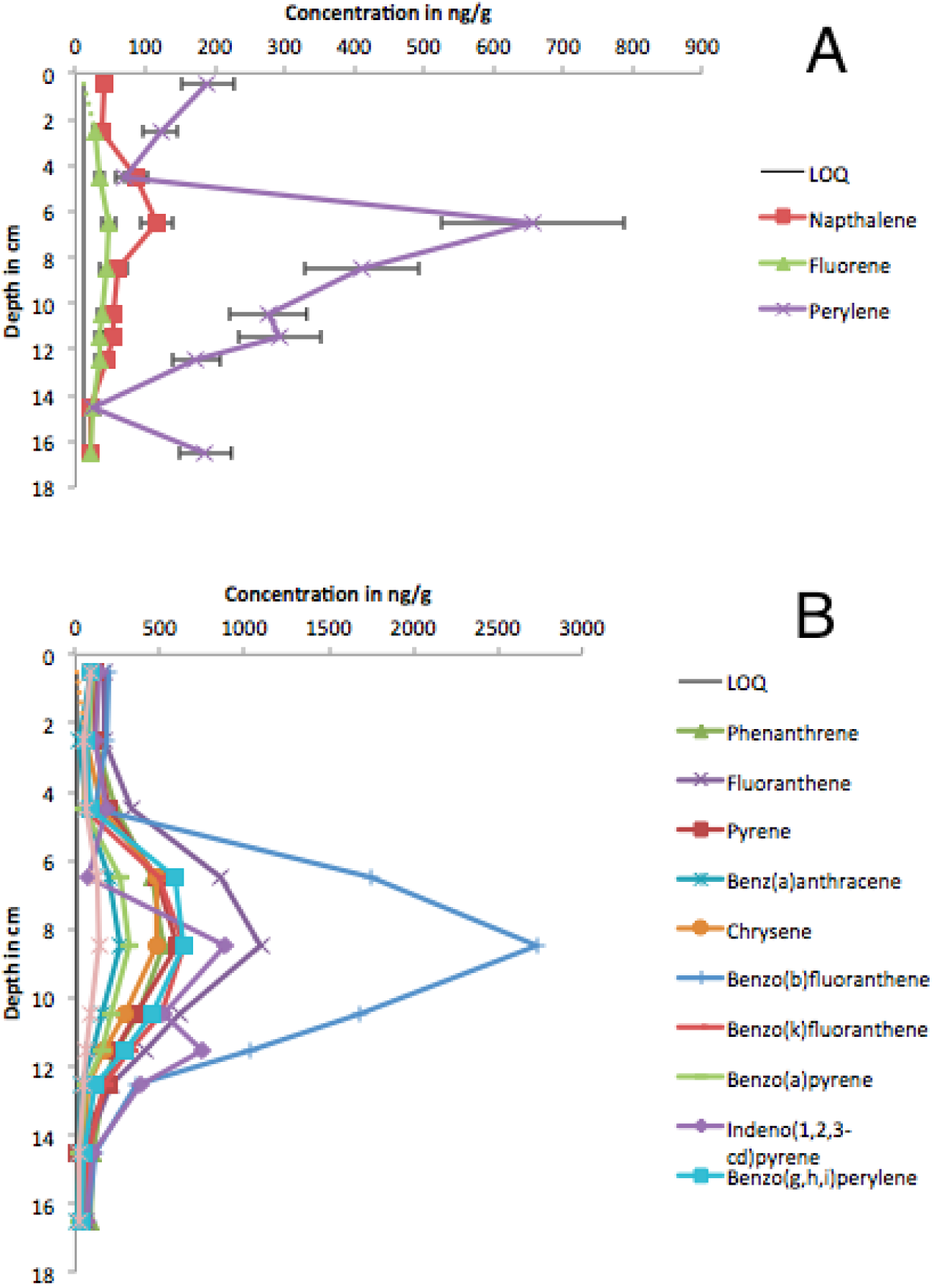
Relative concentrations of quantified PAHs from Solitaire L. A: depicts the aberrant PAHs that lack that general profile pattern. B: depicts the congruent PAHs that follow the general profile pattern with a maximum concentration at 8.5 cm. Error bars represent a 20% relative standard deviation and were omitted from B to improve visibility. The limit of quantification (LOQ) is shown by the vertical black line. The dashed line indicates concentrations falling below the LOQ.

**Figure 4.2.2:**
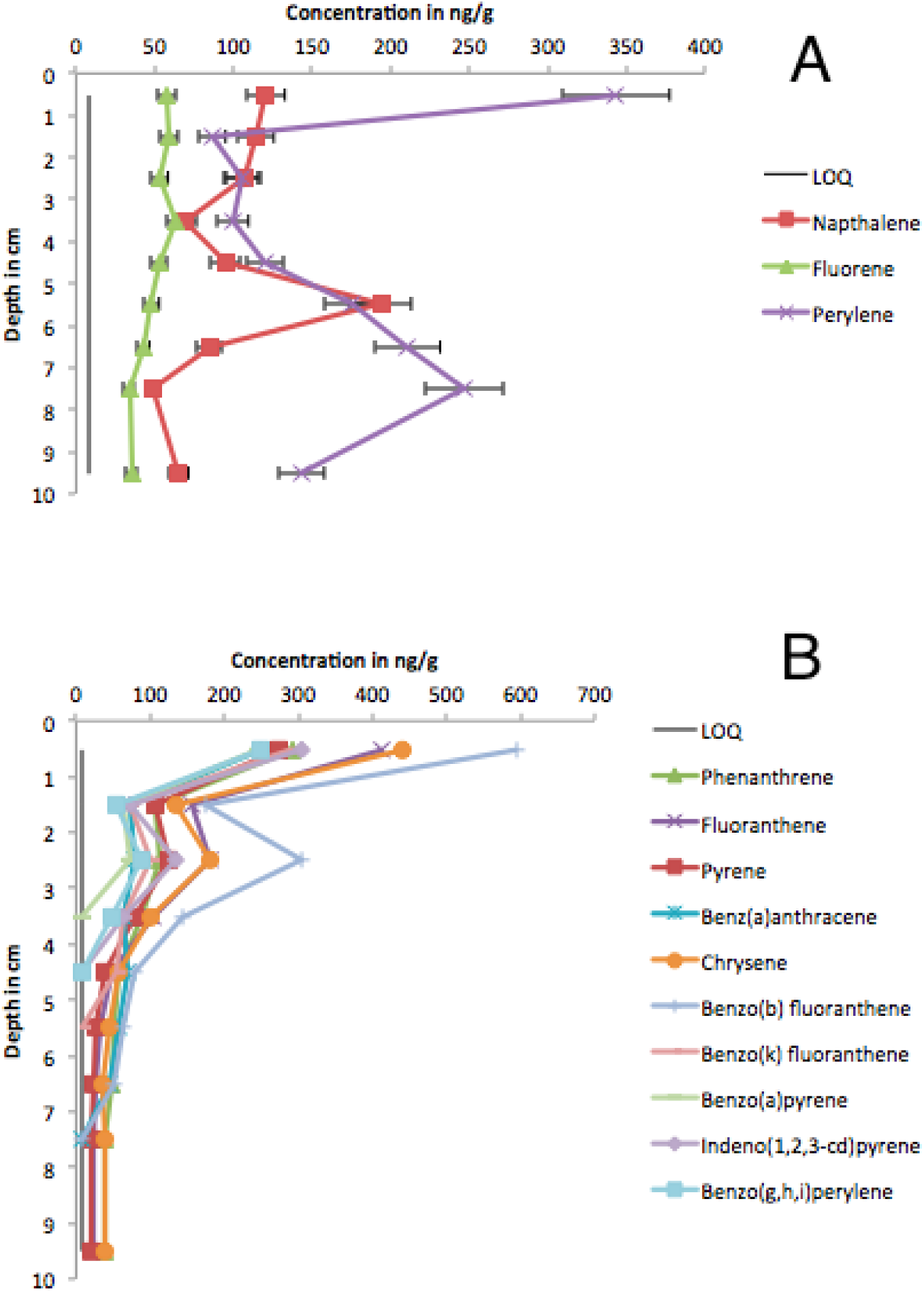
Relative concentrations of quantified PAHs from Fairbank L. A: depicts the aberrant PAHs that lack that general profile pattern. B: depicts the congruent PAHs that follow the general profile pattern with a local maximum concentration at a depth of 2.5 cm. Error bars represent a 10% relative standard deviation and were omitted from B to improve visibility. The vertical black line indicates the limit of quantification (LOQ).

In both profiles the high molecular weight PAHs with four or more aromatic rings were typically found in greater concentrations than those of low molecular weight (Figure 4.3.1). Notably, the dominant pyrogenic PAH in each profile was benzo[b]fluoranthene. Many of the low molecular weight PAHs were either below the level of quantification (BLOQ) or below level of detection (BLOD). All PAHs in Solitaire L. sediments were quantified except for two low molecular weight PAHs, acenapthylene and acenaphtene, which were below the level of detection, and anthracene, which was below the level of quantification and co-eluted with a strongly inferring non-PAH peak. In the Fairbank L. sediments all PAHs were quantified except for: acenaphthylene, acenaphthene, anthracene, and dibenz[a,h]anthracene which were below the limit of detection.

**Figure 4.3.1:**
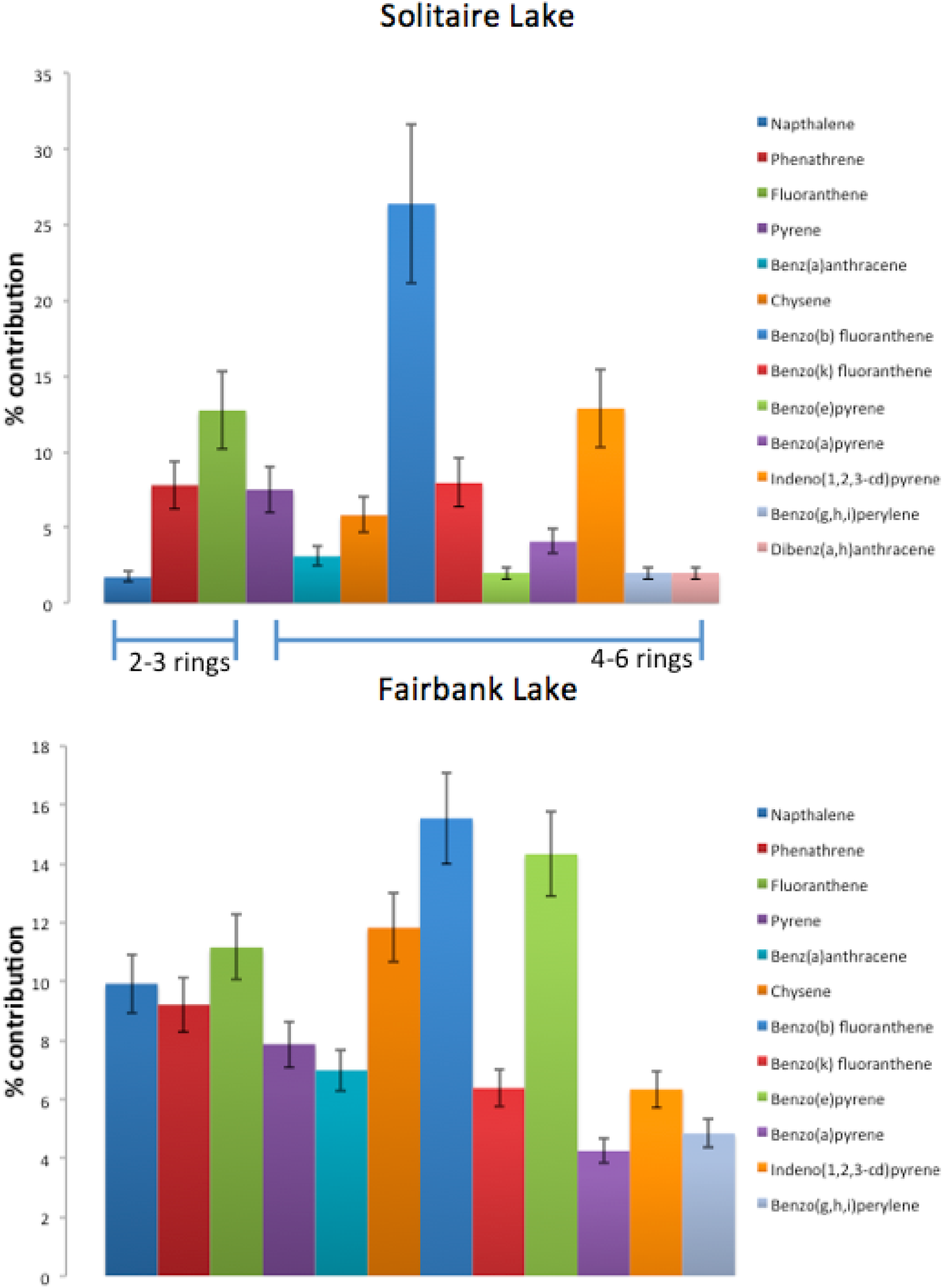
Relative contribution of identified PAHs to total measured pyrogenic PAH concentration. For Solitaire Lake the error bars display a 20% RSD, while for Fairbank Lake the error bars display a 10% RSD based on standard replicates. Note dibenz[a,h]anthracene was below the limit of detection in Fairbank L.

## 5.0 Discussion and Analysis

### 5.1 Pyrogenic PAH Profiles

The PAH profiles observed at the two sites in this study each present distinctive patterns that agree with previous research. The distinctive contaminant pattern between lakes in relatively close proximity (within 200 km) is common (Kannan et al. 2005). The Solitaire L. PAH profile has a similar to trend to those observed in Siskiwit L. first by McVeety and Hites (1988) and again by Slater et al. (2013). The overall shapes of these profiles are consistent, showing decreasing concentrations at the upper and lower ends of the core. Additionally, this data parallels studies from Pettaquamscut River in Rhode Island (Lima et al., 2003) and remote European lakes (Fernandez et al., 2000), which both noted a rise from background levels (20-100 ng/g dw) to maximum peaks (over 1000 ng/g) at the turn of century 1960-1980 (mid profile), before values decreased to present day levels, which were still in some cases 3-20 times higher than background levels. In Solitaire L., the upper profile concentrations were an order of magnitude larger than the background levels. The Fairbanks profile more closely resembled sediment profiles from various locations in the United States, noting increasing PAH concentrations in more recently deposited sediments (Van Metre et al., 2003; Van Metre and Mahler, 2005).

Overall the concentrations from both sites in this study fall within the expected range. Solitaire L. had a maximum concentration of 8438 ng/g dw and Fairbank L. 3500 ng/g dw whereas previous studies of Siskitwit L., Lake Michigan and Pettaquamscutt River had maximum peak values ranging from 900 ng/g dw, 3750 ng/g dw and 9000 ng/g dw, respectively (McVeety and Hites, 1988; Schneider et al., 2001; Lima et al., 2003).

The most significant difference between the two sites aside from the total PAH concentration is the significant increase in PAH deposition that Fairbank L. experiences that is absent in Solitaire L. Assuming that the local maximum in Fairbanks L. and the absolute maximum in Solitaire L. correspond to the same point in time, the historical maximum total PAH concentration in Solitaire L. (8438 ng/g) is greater than Fairbank L. (1592 ng/g) by 530%. This suggests that historically, likely during the 1960-1980s, Solitaire L. was more heavily influenced by atmospheric deposition of pyrogenic PAHs. In the most recent sediments the concentrations in Fairbank L. (3532 ng/g) exceed the concentrations in Solitaire L. (1239 ng/g) by 290%. These recent difference suggest that the Fairbank L. is current receiving more pyrogenic PAHs due to atmospheric deposition compared to its southern counterpart Solitaire L. This shift in trends could be the result of increasing population growth within the Greater Sudbury Region, as similar upward trends in PAH concentrations share a strong association with rapid urbanization and increasing vehicle traffic (Van Metre and Mahler, 2005).

The differences in profiles between Solitaire L. and Fairbanks L. suggest differing extents of influence from various sources. However, in each profile there is a similar dominance of benzo[b]fluoranthene suggesting that there may be an influence from a geographically distinct yet similar source, or there may be an influence from a major regional source. There are many potential PAH sources, although coke ovens, automobile traffic, wood and coal burning are considered the major sources in recent years (Kannan et al. 2005). The relative proportions of the contributing PAHs can further indicate potential sources, as each source can contribute specific profiles or signatures (Khali et al., 1995). Both sites in this study agree with the observations of Kannan et al. (2005), with four- and five-ring PAHs as the most abundant compounds in the profile. At both sites the more recalcitrant larger PAHs contribute to over 70% of the detected PAHs, while the more labile two- and three-ring PAHs (acenaphthylene, acenaphthene, and anthracene) are less abundant and often below the level of detection. These low molecular weight PAHs may be in low abundance because of their low concentration in the major contributing sources, or potentially because of their volatility. Wilcke (2007) observed that larger contributions of high molecular weight compounds are associated with increasing atmospheric PAH deposition. Up to 80% of low molecular weight PAHs such as phenanthrene and anthracene may be lost through surface volatilization before sedimentation (McVeety and Hites, 1988). As vapour pressure decreases with increased molecular weight, less of the high molecular weight PAHs are lost to the atmosphere and greater amounts are deposited in the sediment. Larger 4-6 ring PAHs predominantly found in the particle phase may be transported over long distances, approximately 1300 km prior to deposition (Windsor and Hites, 1979). The dominance of high molecular weight over low molecular weight PAHs in both profiles suggests that both sites are predominantly influenced by atmospheric deposition that is not in the immediate vicinity of an influence point source. Ritcher and Howard (2000) also observed that parent PAHs with 3, 4, and 5 rings dominate with emissions from wood and coal combustion, and vehicle exhaust, implicating these as potential sources to these two lakes.

In the most recent depositional history, Fairbank L. sediment is more contaminated than Solitaire L. This is observed in terms of absolute PAH concentrations, as well as the PAH distribution. A previous study compared two sites in Hamilton Harbour finding that the more contaminated site was in closer proximity to a local contaminant source and low molecular weight PAHs were present in higher concentrations (Morrill et al., 2014). Fairbanks L. had a greater proportion of low molecular weight PAH and shared a conspicuous PAH distribution with those observed in Hamilton Harbor. This may provide evidence that the non-point sources contributing to contamination in Fairbank L. are closer than those contributing to Solitaire L.

PAH distribution is further a useful tool used to distinguish petrogenic and pyrogenic sources. In both studied sites parent PAHs dominate over alkylated PAHs (that were not observed at either site) and high molecular weight PAHs dominate over low molecular weight PAHs providing strong evidence that the sources of these sedimentary PAHs are pyrogenic (Wang and Fingas, 2003). In terms of distinguishing between petroleum and combustion and discriminating amongst potential sources the fluoranthene/(fluoranthene + pyrene) [Fl/(Fl + Py)] using ion mass 202, indeno(1,2,3-cd)pyrene/(indeno(1,2,3-cd)pyrene + benzo(g,h,i)perylene) [IP/(IP + Bghi)] using ion mass 202, indeno(1,2,3-cd)pyrene/(indeno(1,2,3-cd)pyrene + benzo(g,h,i)perylene) [IP/(IP + Bghi)] using ion mass 276, and benzo(a)anthracene/(benzo(a)anthracene + chrysene/triphenylene) [BaA/Σ228] ratios have previously been used (McVeety and Hites, 1998; Yunker et al., 2002; Wang and Fingas, 2003; Morrill et al., 2014). The ratios used are elevated in both locations with minimal variation when depth changes implying combustive sources have had a continuous depositional influence in Ontario (Table 5.1.1).

**Table 5.1.1:** Whole core PAH ratios for Solitaire L. and Fairbank L.

| Solitaire lake |  |  |  |  |  |  |  |  |  |  |  |
| --- | --- | --- | --- | --- | --- | --- | --- | --- | --- | --- | --- |
| Depth (cm) | 0-1 | 2-3 | 4-5 | 5-6 | 6-7 | 8-9 | 10-11 | 12-13 | 14-15 | 16-17 | Mean $\pm SD$ |
| Fl/(Fl + Py) | 0.60 | 0.60 | 0.62 | 0.64 | 0.64 | 0.64 | 0.64 | 0.51 | 0.84 | 0.64 | $0.64 \pm 0.08$ |
| IP/(IP + Bghi) | 0.62 | 0.65 | 0.66 | 0.67 | 0.58 | 0.53 | 0.72 | 0.77 | 0.70 | 0.67 | $0.60 \pm 0.07$ |
| BaA/$\Sigma 228$ | n.d. | 0.28 | 0.27 | 0.30 | 0.35 | 0.34 | 0.37 | 0.35 | 0.28 | 0.28 | $0.31 \pm 0.04$ |
| Fairbank lake |  |  |  |  |  |  |  |  |  |  |  |
| Depth (cm) | 0-1 | 1-2 | 2-3 | 3-4 | 4-5 | 5-6 | 6-7 | 7-8 | 9-10 | Mean $\pm SD$ | |
| Fl/(Fl + Py) | 0.60 | 0.59 | 0.59 | 0.57 | 0.55 | 0.55 | 0.55 | 0.54 | 0.54 | $0.57 \pm 0.02$ | |
| IP/(IP + Bghi) | 0.55 | 0.57 | 0.61 | 0.57 | n.d. | n.d. | n.d. | n.d. | n.d. | $0.58 \pm 0.02$ | |
| BaA/$\Sigma 228$ | 0.36 | 0.35 | 0.30 | 0.39 | 0.55 | 0.56 | 0.55 | n.d. | n.d. | $0.44 \pm 0.11$ | |

n.d. = no data
SD = standard deviation

Using the whole core average the Fl/(Fl + Py) ratio was calculated to be 0.64±0.08 in Solitaire L. and 0.57±0.02 in Fairbank L., and the IP/(IP + Bghi) was calculated to be 0.60±0.07 in Solitaire L. and 0.58±0.02 in Fairbank L. For both Fl/(Fl + Py) and IP/(IP + Bghi), low ratios of (<40 and <20, respectively) indicate non-combustive petroleum; intermediate ratios of (0.40-0.50 and 0.20-0.50, respectively) indicate combustion of liquid fossil fuel (gasoline, diesel and crude oil); and high ratios >0.50 are characteristic of grass, wood and coal combustion (Yunker et al., 2002; Yunker and MacDonald, 2003). The BaAΣ228 ratios in Solitaire L. and Fairbank L. were greater than 0.2 indicating combustion sources. Both sites observed have diagnostic ratios above 0.50 for Fl/(Fl + Py) and IP/(IP + Bghi) and above 0.2 for BaA/Σ228 indicating that the combustion of grass, wood and coal are the predominant inputs in these regions. The ratios from Fairbank L. and Solitaire L. plot most closely to wood, coal and gasoline combusted sources (Figure 5.1.1) when comparing IP/(IP + Bghi) to BaA/Σ228 for other combusted and non-combusted fossil fuels.

**Figure 5.1.1:**
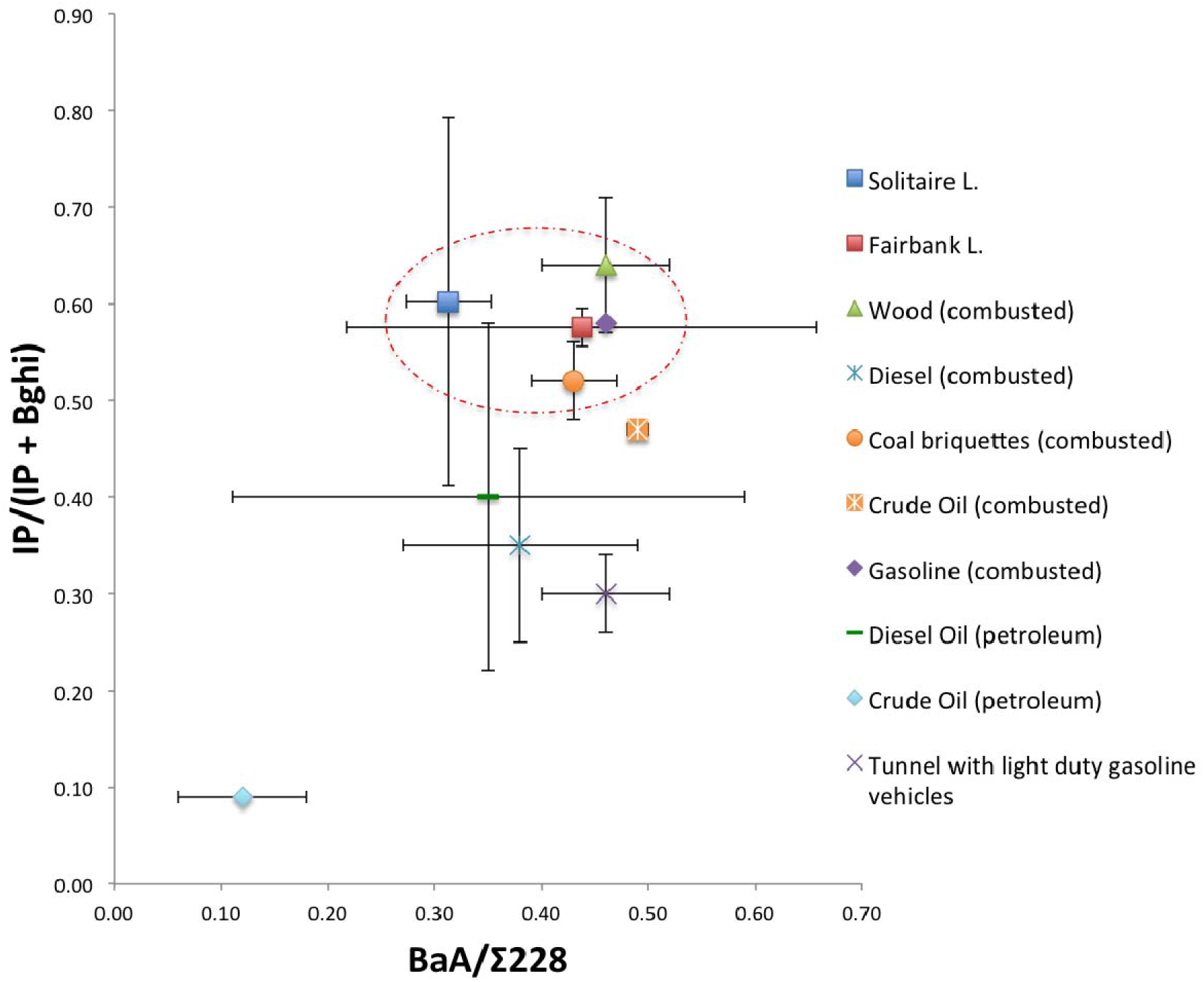
PAH ratios for Solitaire L. and Fairbank L., plotted with diagnostic ratios for IP/(IP + Bghi) versus BaA/ 228 for combusted and non-combusted fuels from Yunker et al. (2002).

Considering the main pyrogenic PAH sources, coke oven and automobile emissions have a low probability as a major potential source because they mostly produce low molecular weight PAHs that were not abundant at these sites (Khalili et al., 1995; Yang et. al, 1998). In both sites, the PAH distribution implicates a non-mobile source as the main source contributor of PAH contaminants, most likely historical coal combustion from power generation or wood combustion from wildfires and heating. However, automobile emissions inevitably have some contribution because of their frequent usage. Based on unpublished trace metal data from the same sites, smelting activities could play a significant role in the deposition of the PAHs found in these profiles (Cheyne, unpublished). However, with the data provided in this study alone a definitive source cannot be identified as each suspected source has non-unique relative proportions of PAH species. Additionally, the relative proportions of the species are not highly conserved between emission source and point of measurement due to volatilization and photodegradation (Lima et al, 2005).

The PAH profiles also indicated the presence of benzo[e]pyrene, which is not included in Restek SV Calibration Mix #5. Benzo[e]pyrene was identified based on the ion extracted, the mass spectrum and comparable elution time based on Agilent Technologies’ “Essential Chromatography and Spectroscopy Catalog 2009-2010 Edition” for PAHs. The presence of this pyrogenic PAH has been noted in previous studies (McVeety and Hites, 1998; Van Metre et al., 2000). Benzo[e]pyrene is often indicative of combustion and is associated with automobile exhaust in high atmospheric concentrations on urban streets (Nielsen et al., 1996). The presence of benzo[e]pyrene particularly in Fairbank L. may indicate the potential influence of traffic related combustion sources.

### 5.2 Perylene

In many studies perylene has been noted to have a distinct behavior from pyrogenic PAHs. Perylene has been the focus of a number of studies because of the profiles of increasing concentrations with depth, indicating it is not carried and deposited in the sediment, but formed after deposition (Gschwend et al., 1981; Silliman et al., 1998; Grice et al., 2009; Slater et al., 2013). The leading hypothesis suggests the majority of observed perylene in these profiles is the result of in situ production by microbial communities living in anoxic conditions, rather than anthropogenic production (Gschwend et al., 1981; Vankatesan, 1988). Undoubtedly some perylene observed at these sites is still the result of atmospheric particulate deposition, yet in this study perylene was omitted from total PAH concentration as it deviates from the definitive pyrogenic PAH trend.

In Solitaire L., the maximum perylene concentration occurs at 6.5 cm (2 cm above the pyrogenic PAH peak), only to decline rapidly before increasing again at 14.5 cm (Figure 5.2.1). Similar observations have been made regarding perylene concentrations in Lake Ontario where they are lower at the surface, peak earlier than pyrogenic PAHs and remain elevated down core (Silliman et al., 1998). At the depth where the maximum perylene concentration of 650 ng/g occurs, it is a dominant compound in the profile exceeding all other PAHs concentrations except for benzo[b]fluoranthene. At the lower depth of 16.5 cm the perylene concentration exceeds all other PAHs. This increasing trend possibly continues at lower depths, however samples were only analyzed to 16.5 cm.

**Figure 5.2.1:**
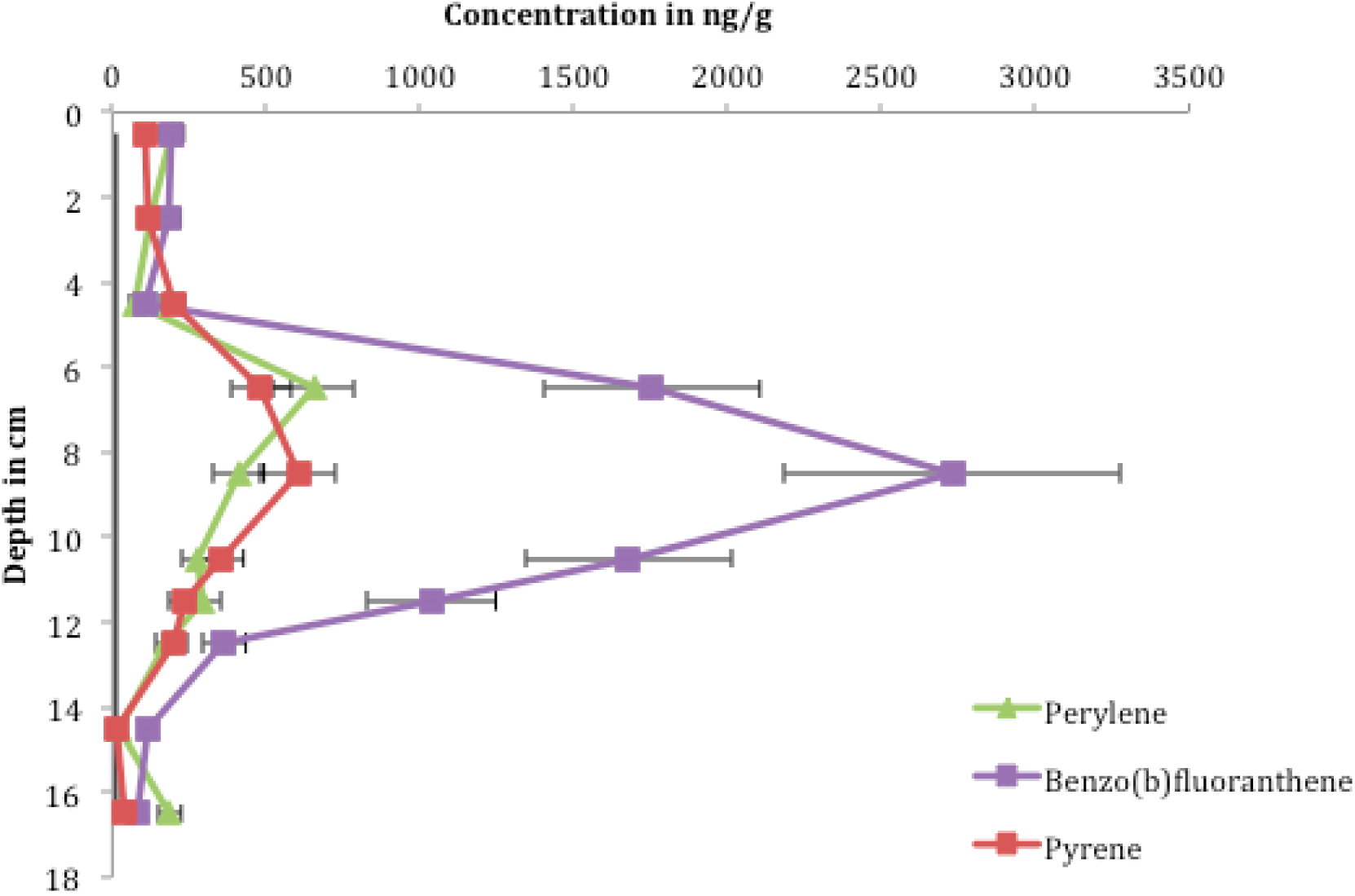
Comparison of perylene to pyrogenic PAH deposition in Solitaire L. The limit of quantification (LOQ) is shown by the vertical black line.

In Fairbank L., the perylene profile correlates with the perylene peaks observed by McVeety and Hites (1988) and Slater et al. (2013). Notably, when the perylene profile reaches its relative maximum concentration, at 247 ng/g, all other pyrogenic PAHs are below background levels at 50 ng/g (Figure 5.2.2). However, at the top of the core perylene concentration increases coincident with the increasing pyrogenic PAHs, reaching a maximum concentration of 340 ng/g. The relatively large abundance of perylene in the uppermost sediment layer of Fairbank L. is an unexpected observation, as perylene is typically absent or present only in minor concentrations in oxic surface sediments (Silliman et al., 1998). However, Hites et al. (1988) observed perylene concentrations exceeding all other PAHs in the upper core sediments from the Pettaquamscutt River. This may suggest an increase in microbial activity in the upper sediments or a recent local source that produces a high abundance of perylene.

**Figure 5.2.2:**
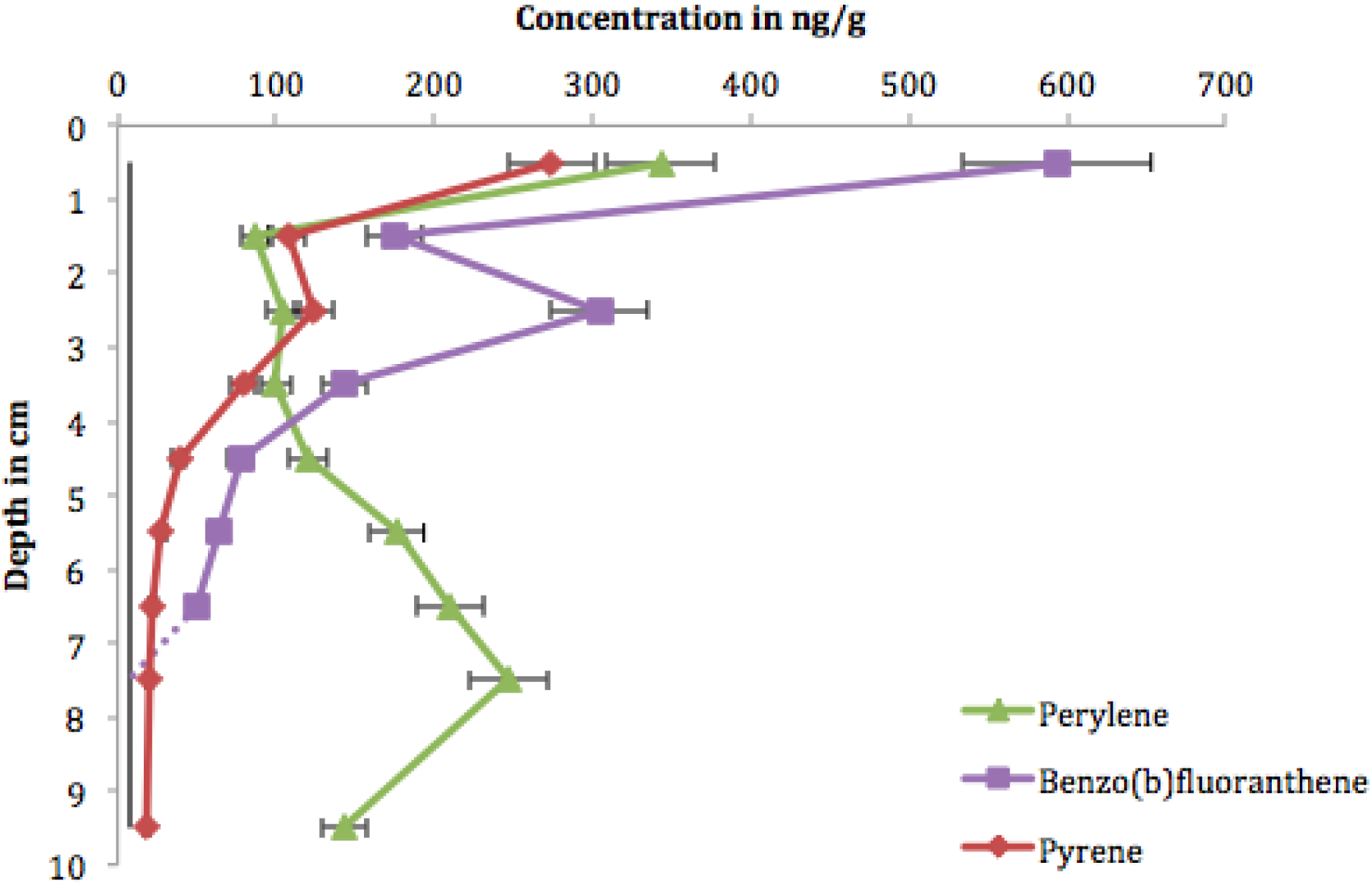
Comparison of perylene to pyrogenic PAH deposition in Fairbank L. The limit of quantification (LOQ) is shown by the vertical black line. The dashed line indicates a compound decreasing below the level of quantification.

The constant exponential trend of increasing perylene concentration with sediment depth observed by Grice et al. (2009) was not verified at either site in this study. In both studied sites, perylene has a distinctive pattern that involved rapid increasing and decreasing sediment fluxes. These perylene fluxes are unlike the pyrogenic PAHs observed, removing the possibility of perylene deposition from the same anthropogenic sources.

### 5.3 Health effects

PAH concentrations in the deposits in the most recent sediments from both lakes exceed the interim sediment quality guidelines (ISQG) established by the Canadian Council of Minister of the Environment (Table 5.3.1). The ISQG values are derived from available scientific information on biological effects of sediment-associated chemicals, and correspond to the threshold level below which adverse biological effects are not anticipated. Benthic organisms that are exposed to high concentrations of PAHs are particularly vulnerable (Van Metre and Mahler, 2005). The PAHs identified in this study are bioavailable and these non-polar compounds can bioaccumulate in the fat and lipid stores of marine invertebrates. The lipid and organic carbon content influence PAH behaviour in water, sediment and tissue, and the octanol-water partitioning coefficient is the single best predictor to determine bioavailability (Douben, 2003). The concentrations of benzo[a]pyrene at each study site well exceed the sediment quality guidelines and should be of particular concern because of carcinogenic properties. Bottom-dwelling fish have demonstrated the growth of malignant tumors after injection of sediments containing PAHs, and particularly high concentrations of benzo(a)pyrene (Metcalfe et al., 1988).

**Table 5.3.1:**
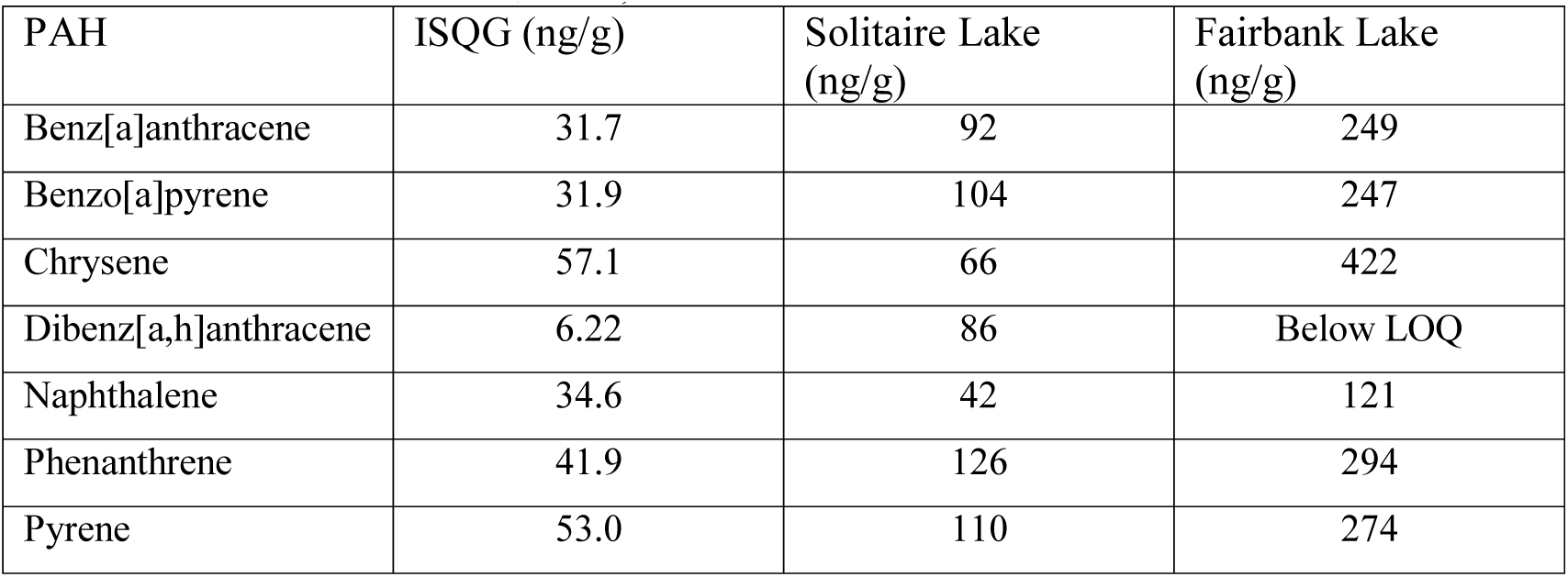
Uppermost sediment concentrations in relation to the ISQG (Canadian Council of Minister of the Environment, 2015)

### 5.4 Method Remarks

As noted above, the corrected concentration values based on the recoveries for the external standards m-terphenyl and 9,10-dihydrophenathracene were used for further comparisons. The precedent to use corrected values has been established in other publications (McVeety and Hites, 1988; Slater et al., 2013). In this study the correction with 9,10-dihydrophenathracene was always proportionally greater than the correction with m-terphenyl. This is likely because of the greater volatility of the low molecular weight standard. Although this correction introduces further uncertainty, as not all PAHs will have the same mass losses as these two standards, the corrections do not change the overall depositional trends. As seen in total PAH profiles, correcting concentrations using the recovery standards does not change the observed trend, rather it adjusts the absolute concentration values, for the most part to larger values. In Fairbank L., the corrected values had no impact on the individual PAH profiles; however, in Solitaire L., the maximum peak value became ambiguous for chrysene (Figure 5.4.1). In the chrysene profile from Solitaire L. the corrected values change the depth at which the maximum PAH concentration occurred, altering the observed depositional pattern.

**Figure 5.4.1:**
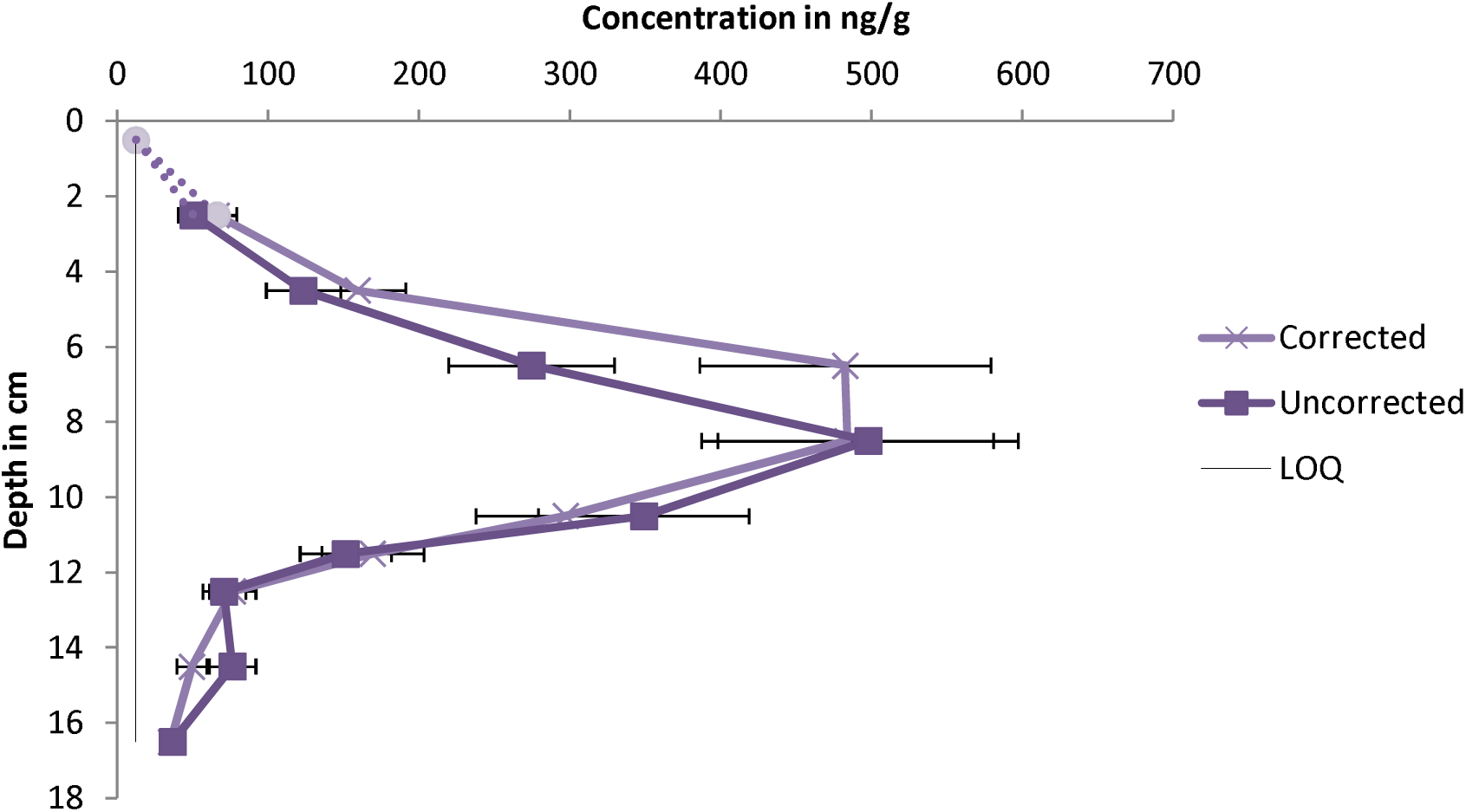
Uncorrected and corrected chrysene profile for Solitaire L. Error bars indicate a 20% RSD and the dashed line represents a concentration BLOQ.

The variance that was provided on each sediment sample was based on the calculated average and standard deviations for o-terphenyl standard injections. Completing this experiment again, at least two triplicate samples should be analyzed, per core, and variances should be calculated per PAH. A per PAH variance will provide a more realistic error as low molecular weight PAHs likely have significantly larger variances based on volatility.

Additionally, the inclusion of a benzo(a)pyrene standard would also benefit the analysis if further research is completed. This would allow for the definitive verification of this compound in the samples, and potential incorporation into the total PAH concentration.

## 6.0 Conclusion

### 6.1 Thesis Summary

Polycyclic aromatic hydrocarbons are mainly produced through the incomplete combustion of organic material (Ravindra et al., 2008a). Their persistence and carcinogenicity present a threat to human and environmental health (Yan et al., 2003). This study confirmed the presence of nearly all 16 priority PAHs in addition to the suspected presence of benzo[e]pyrene in both central Ontario sites. Although there has been a successful abatement of point sources, non-point sources such as atmospheric deposition and surface runoff constitute the major inputs into the environment. These complex PAH assemblages in recent sediments at the two studied sites have geochemical and environmental significance. The dominance of high molecular weight, and unsubstituted PAH content at the studied sites indicate the predominant source is pyrogenic. Overall, the concentrations from both sites in this study fall within the expected range of 900-9000 ng/g dw. The most significant difference between the two sites aside from the total PAH concentration, is the increase in PAH deposition that occurs in Fairbank L. is absent in Solitaire L. The differences in profiles between these sites suggest differing extents of influence from various sources, with each likely having the unique sources for the majority of PAH deposition. This study confirmed that the presence of perylene is likely from non-pyrogenic deposition and potentially in-situ production by microbial communities. The most recent sediments at both locations exceed the Environment Canada sediment quality guidelines, indicating a hazard for aquatic organisms, with broader implications affecting both environmental and human health. This limited study provides meaningful insight regarding the differential atmospheric deposition in central Ontario. In comparison with previous studies by McVeety and Hites (1988), Van Metre and Mahler (2005), and Slater et al. (2013), a distinctive pattern of deposition in each lake is observed. This would suggest that local sources are larger contributors to deposition than a regional influencing source.

### 6.2 Future Directions

This study analyzed pyrogenic PAH deposition in two lakes from central Ontario. An expanded study should include the analysis of more locations that could be influenced by similar regional sources. Due to time requirements, cores that were collected from Semiwite Lake and Big McDougal Lake were unable to be processed. Additionally, this study relied only on depth and PAH concentration to construct profiles. This provides an estimate of chronological events; however, no conclusions on the precise timeframe of deposition can be made without further studies involving lead-210 dating. With the use of lead-210 dating accounting for different deposition rates based on lake morphology, the PAH profiles provide a more rigorous model of atmospheric deposition, which can be more accurately compared across studies using similar dating techniques. Once sediment dating has been completed the profiles can be adjusted to access the changes in PAH levels in terms of flux that are a function of source strength, sedimentation rate and sediment dilution (Van Metre et al., 2000). Trace metal analysis may aid in identifying the potential sources of contamination and can confirm the trends observed in the PAH profiles and the timeframe provided by lead-210 dating. The addition of sedimentation rate and metals data establishes a more meaningful profile, and can allow source attribution prompting the development of strategies for effectively reducing or mitigating the effects of atmospheric PAH deposition.

## Appendix

## Appendix A

**Table 1:** Physical and chemical properties for select PAHs (McKay et al., 1998)

| Compound | No. Rings | MW | S (mg/L) | VP (Pa) | Log Kow |
| --- | --- | --- | --- | --- | --- |
| Napthalene | 2 | 128 | 31 | 10.4 | 3.37 |
| Acenaphthylene | 3 | 150 | 16.1 | 0.9 | 4.00 |
| Acenaphthene | 3 | 154 | 2.1 | 0.3 | 3.92 |
| Fluorene | 3 | 166 | 1.9 | 0.09 | 4.18 |
| Phenanthrene | 3 | 178 | 1.1 | 0.02 | 4.57 |
| Anthracene | 3 | 178 | 0.045 | 0.001 | 4.54 |
| Fluoranthene | 4 | 202 | 0.26 | 0.00123 | 5.22 |
| Pyrene | 4 | 202 | 0.132 | 0.0006 | 5.18 |
| Benz[a]anthracene | 4 | 228 | 0.011 | 2.80E-05 | 5.91 |
| Chrysene | 4 | 228 |  | 5.70E-07 | 5.86 |
| Benzo[b]flouranthene | 5 | 252 | 0.0015 |  | 5.80 |
| Benzo[k]flouranthene | 5 | 252 | 0.0008 | 5.20E-08 | 6.00 |
| Benzo[a]pyrene | 5 | 252 | 0.0038 | 7.00E-07 | 6.04 |
| Perylene | 5 | 252 | 0.0004 | 1.40E-08 | 6.25 |
| Indeno[1,2,3-cd]pyrene | 6 | 278 |  |  |  |
| Dibenz[a,h]anthracene | 5 | 278 | 0.0006 | 3.70E-10 | 6.75 |
| Benzo[g,h,i]perylene | 6 | 268 | 0.00026 |  | 6.50 |

MW= molecular weight; S= solubility in water; VP= vapour pressure; Kow= octanol-water partitioning coefficient.

## Appendix B

**Figure 1:**
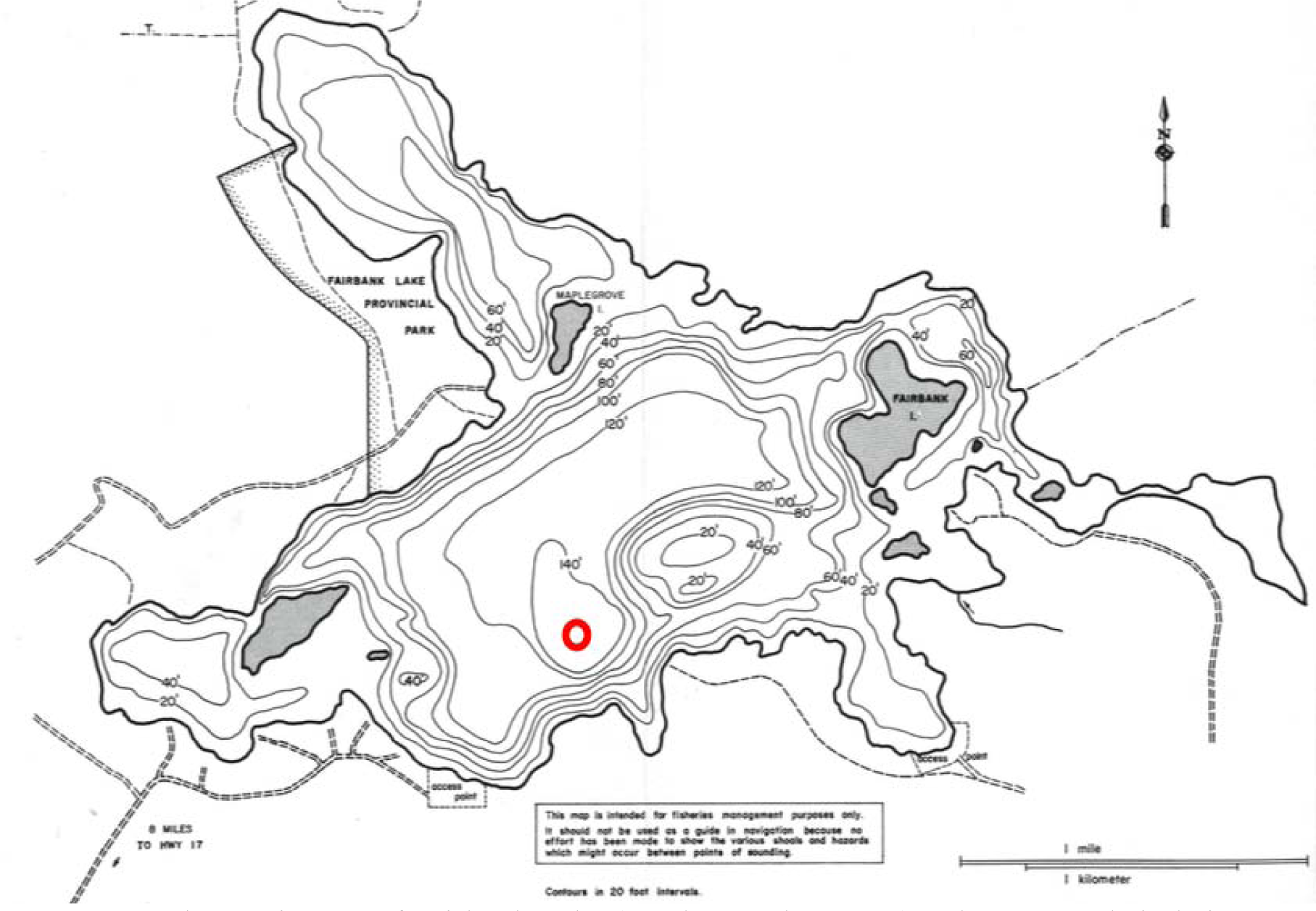
Bathymetric map of Fairbank Lake (Angler’s Atlas, 2015). The open-red circle is an approximation of the sampling location (DMS: 46°27’33.5N, 81°25’53.3W).

**Figure 2A:**
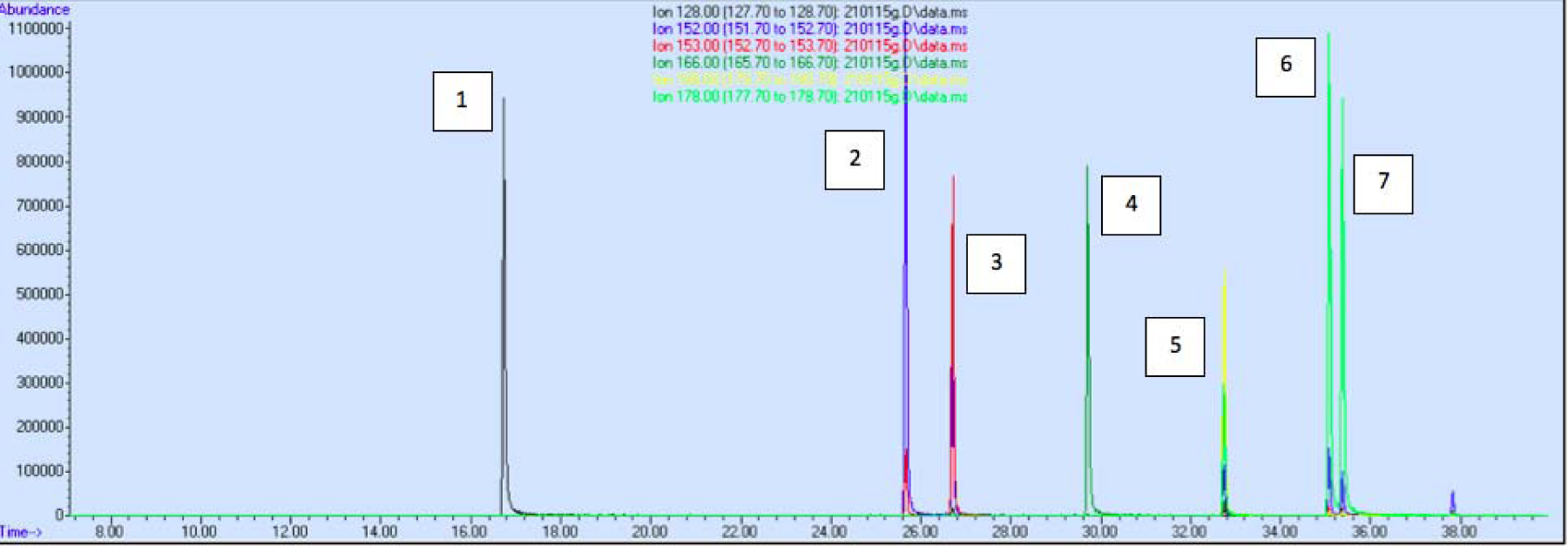
Extracted ion chromatogram of LWM PAH standards. The peaks in this 10 ppm standard are: 1) napthalene; 2) acenapthylene; 3) acenapthene; 4) fluorene; 5) 9,10-dihydrophenanthrene; 6) phenanthrene; 7) anthracene.

**Figure 2B:**
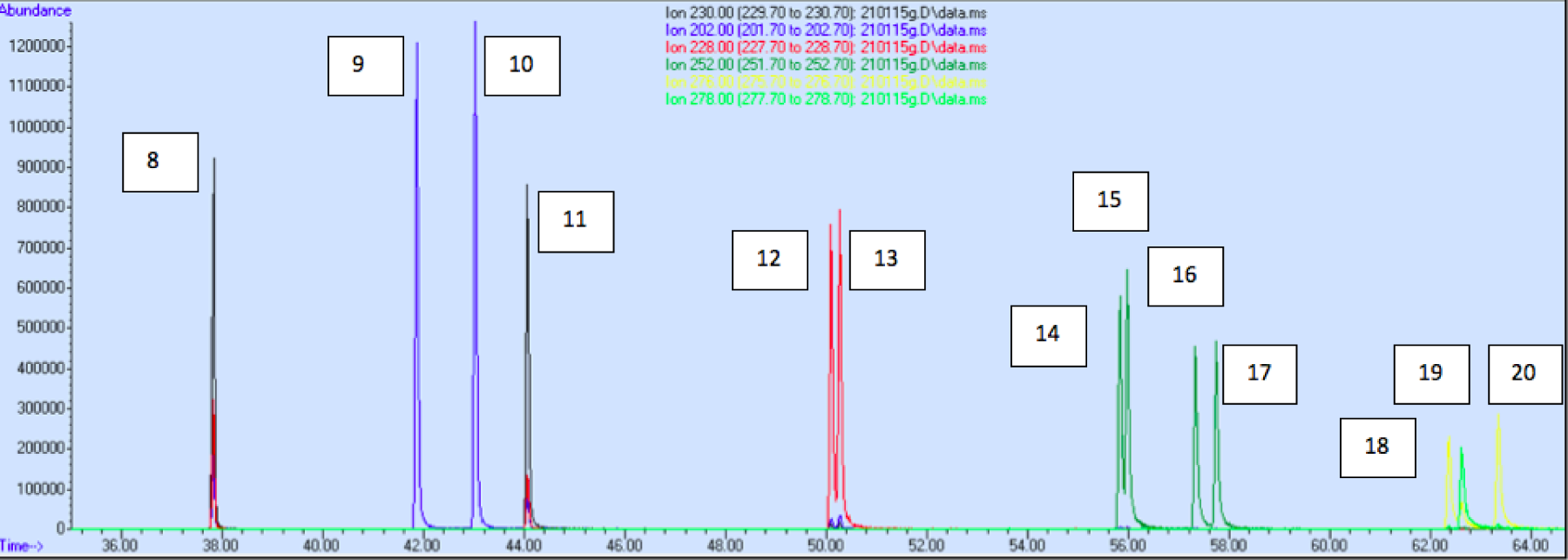
Extracted ion chromatogram of the HMW PAH standards. The peaks in this 10 ppm standard are: 8) o-terphenyl; 9) fluoranthene; 10) pyrene; 11) m-terphenyl; 12) benz[a]anthracene; 13) chrysene; 14) benzo[b]flouranthene; 15) benzo[k]flouranthene; 16) benzo[a]pyrene; 17) perylene; 18) indeno[1,2,3-cd]pyrene; 19) dibenz[a,h]anthracene; 20) benzo[g,h,i]perylene.

**Table B1A:** Wet and dry soil weights and density measurements for Solitaire L. core F.

| Solitaire Core F |  |  |  |  |  |  |  |  |
| --- | --- | --- | --- | --- | --- | --- | --- | --- |
| Core section cm | Wet mass soil+ bag(g) | Wet mass soil(g) | Dry mass soil + bag (g) | Dry mass soil(g) | % Soil Weight | % Water | Wet Bulk Density (g/cm<sup>3</sup>) | Dry Bulk Density (g/cm<sup>3</sup>) |
| 0-1 | 27.6214 | 24.28 | 4.2497 | 0.90 | 15 | 85 | 0.73 | 0.03 |
| 2-3 | 28.8441 | 25.50 | 4.5537 | 1.21 | 16 | 84 | 0.77 | 0.04 |
| 4-5 | 41.4718 | 38.13 | 4.8310 | 1.49 | 12 | 88 | 1.15 | 0.04 |
| 6-7 | 39.4363 | 36.09 | 5.3634 | 2.02 | 14 | 86 | 1.09 | 0.06 |
| 8-9 | 28.9753 | 25.63 | 5.6345 | 2.29 | 19 | 81 | 0.77 | 0.07 |
| 10-11 | 35.5820 | 32.24 | 5.6767 | 2.33 | 16 | 84 | 0.97 | 0.07 |
| 11-12 | 38.2277 | 34.88 | 6.0321 | 2.69 | 16 | 84 | 1.05 | 0.08 |
| 12-13 | 31.6434 | 28.30 | 4.3904 | 1.04 | 14 | 86 | 0.85 | 0.03 |
| 14-15 | 35.3669 | 33.37 | 5.7521 | 3.75 | 16 | 84 | 1.01 | 0.11 |
| 16-17 | 38.0000 | 36.00 | 4.5039 | 2.50 | 12 | 88 | 1.08 | 0.08 |

**Table B1B:** Wet and dry soil weights and density measurements for Solitaire L. core E.

| Solitaire Core E |  |  |  |  |  |  |  |  |
| --- | --- | --- | --- | --- | --- | --- | --- | --- |
| Core section<br>cm | Wet mass<br>soil+<br>bag(g) | Wet mass<br>soil(g) | Dry mass<br>soil + bag<br>(g) | Dry mass<br>soil(g) | % Soil<br>Weight | % Water | Wet Bulk<br>Density<br>(g/cm<sup>3</sup>) | Dry Bulk<br>Density<br>(g/cm<sup>3</sup>) |
| 0-1 | 36.73 | 33.39 | 4.12 | 0.77 | 11 | 89 | 1.006 | 0.02 |
| 2-3 | 35.29 | 31.94 | 4.57 | 1.22 | 13 | 87 | 0.963 | 0.04 |
| 4-5 | 36.28 | 32.93 | 4.96 | 1.61 | 14 | 86 | 0.993 | 0.05 |
| 6-7 | 35.48 | 32.13 | 5.52 | 2.17 | 16 | 84 | 0.968 | 0.07 |
| 8-9 | 36.82 | 33.47 | 5.75 | 2.40 | 16 | 84 | 1.009 | 0.07 |
| 10-11 | 39.39 | 36.04 | 5.53 | 2.18 | 14 | 86 | 1.086 | 0.07 |
| 11-12 | 34.73 | 31.38 | 5.92 | 2.57 | 17 | 83 | 0.946 | 0.08 |
| 12-13 | 40.73 | 37.38 | 6.15 | 2.80 | 15 | 85 | 1.127 | 0.08 |
| 14-15 | 36.33 | 32.98 | 5.96 | 2.61 | 16 | 84 | 0.994 | 0.08 |
| 16-17 | 35.96 | 32.61 | 6.10 | 2.75 | 17 | 83 | 0.983 | 0.08 |

**Table B1C:** Wet and dry soil weights and density measurements for Fairbanks L. core D.

| Fairbank Core D |  |  |  |  |  |  |  |  |
| --- | --- | --- | --- | --- | --- | --- | --- | --- |
| Core section<br>cm | Wet mass<br>soil+<br>bag(g) | Wet mass<br>soil(g) | Dry mass<br>soil + bag<br>(g) | Dry mass<br>soil(g) | % Soil<br>Weight | % Water | Wet Bulk<br>Density<br>(g/cm<sup>3</sup>) | Dry Bulk<br>Density<br>(g/cm<sup>3</sup>) |
| 0-1 | 51.5555 | 49.17 | 6.6949 | 4.31 | 13 | 87 | 1.48 | 0.13 |
| 1-2 | 37.8628 | 35.48 | 7.2604 | 4.88 | 19 | 81 | 1.07 | 0.15 |
| 2-3 | 39.1980 | 36.82 | 8.0189 | 5.64 | 20 | 80 | 1.11 | 0.17 |
| 3-4 | 42.0930 | 39.71 | 8.5217 | 6.14 | 20 | 80 | 1.20 | 0.19 |
| 4-5 | 36.3580 | 33.98 | 7.7316 | 5.35 | 21 | 79 | 1.02 | 0.16 |
| 5-6 | 38.5571 | 36.18 | 8.3311 | 5.95 | 21 | 79 | 1.09 | 0.18 |
| 6-7 | 40.2131 | 37.83 | 9.2960 | 6.91 | 23 | 77 | 1.14 | 0.21 |
| 7-8 | 39.4202 | 37.04 | 9.4064 | 7.02 | 23 | 76 | 1.12 | 0.21 |
| 9-10 | 41.7912 | 39.41 | 10.1325 | 7.75 | 24 | 76 | 1.19 | 0.23 |

**Table B2:** Quantification ions and retention times used for GC/MS analysis

| PAH | Quantifying Ion | Retention Time |
| --- | --- | --- |
| Napthalene | 128 | 16.736 |
| Acenaphthylene | 152 | 25.662 |
| Acenaphthene | 153 | 26.708 |
| Fluorene | 166 | 29.710 |
| 9,10-dihydrophenanthrene | 180 | 32.743 |
| Phenanthrene | 178 | 35.079 |
| Anthracene | 178 | 35.377 |
| o-terphenyl | 230 | 37.827 |
| Fluoranthene | 202 | 41.824 |
| Pyrene | 202 | 43.029 |
| m-terphenyl | 230 | 44.064 |
| Benz[a]anthracene | 228 | 50.092 |
| Chrysene | 252 | 50.267 |
| Benzo[b]flouranthene | 252 | 55.829 |
| Benzo[k]flouranthene | 252 | 55.977 |
| Benzo[a]pyrene | 252 | 57.330 |
| Perylene | 252 | 57.744 |
| Indeno[1,2,3-cd]pyrene | 276 | 62.361 |
| Dibenz[a,h]anthracene | 278 | 62.622 |
| Benzo[g,h,i]perylene | 276 | 63.341 |

